# Long-read genome assembly of the insect model organism *Tribolium castaneum* reveals spread of satellite DNA in gene-rich regions by recurrent burst events

**DOI:** 10.1101/2023.04.24.538079

**Authors:** Volarić Marin, Despot-Slade Evelin, Veseljak Damira, Mravinac Brankica, Meštrović Nevenka

## Abstract

Eukaryotic genomes are replete with satellite DNAs (satDNAs), large stretches of tandemly repeated sequences which are mostly underrepresented in genome assemblies. Here we combined Nanopore long-read sequencing with a reference-guided assembly approach, to generate an improved, high-quality genome assembly TcasONT of the model beetle *Tribolium castaneum*. Enriched by 45 Mb in the repetitive part, the new assembly comprises almost the entire genome sequence. We used the enhanced assembly to conduct global and in-depth analyses of abundant euchromatic satDNAs, Cast1-Cast9. Contrary to the commonly adopted view, our finding showed the extensive spread of satDNAs in gene-rich regions, including long arrays. The results of the principal component analysis of monomers and sequence similarity relationships between satDNA arrays, revealed an occurrence of recent satDNAs array exchange between different chromosomes. We proposed a scenario of their genome dynamics characterized by repeated bursts of satDNAs spreading through euchromatin, followed by a process of elongation and homogenization of arrays. We also found that suppressed recombination on the X chromosome has no significant effect on the spread of satDNAs, but rather tolerates the amplification of these satDNAs into longer arrays. Analyses of arrays’ neighboring regions showed a tendency of one Cast satDNA to be associated with transposon-like elements. Using 2D electrophoresis followed by Southern blotting, we proved Cast satDNAs presence in the fraction of extrachromosomal circular DNA (eccDNA). We point to two mechanisms that enable the said satDNA spread to occur: transposition by transposable elements and insertion mediated by eccDNA. The presence of such a large proportion of satDNA in gene-rich regions inevitably gives rise to speculation about their possible influence on gene expression.

## Background

Satellite DNAs (satDNAs) are the most abundant and rapidly evolving non-coding DNA of all eukaryotic genomes. In contrast to other groups of repetitive elements such as interspersed transposable elements, which are scattered throughout the genome, satDNAs are characterized by monomer sequences tandemly arranged into long arrays. Previous studies on many animal and plant species indicate preferential localization of satDNAs in the (peri)centromeric heterochromatin [1]. Heterochromatin represents a highly compact gene-poor specialized chromatin structure that is distinct from the gene-rich euchromatin. SatDNAs in (peri)centromeric heterochromatin have a long history of investigation including many plant and animal species, and studies have suggested a role of satDNA in chromatin packaging, centromere formation/maintenance, and centromere evolution [2–4]. SatDNAs are also linked to chromosome mis-segregation [3], disease phenotypes [5], and reproductive isolation between species [6]. In addition to standard satDNA-rich regions such as (peri)centromeric heterochromatin, recent studies revealed examples where short arrays of (peri)centromeric satDNAs also spread to euchromatic regions. For example, (peri)centromeric satDNA families in humans, *Drosophila* and the beetle *Tribolium castaneum* can also be dispersed within euchromatin in the form of clustered or short repeat arrays [7–9]. In *Drosophila,* X-specific (peri)centromeric 1688 satDNA repeats dispersed in euchromatic regions have a role in dosage compensation through up-regulation of X-linked genes in *Drosophila* males [10,11]. The euchromatic counterpart of (peri)centromeric DNA in the beetle *T. castaneum* modulates the local chromatin environment upon heat stress changing the expression of neighboring genes [12]. Although there is clear evidence that satDNAs exist outside the (peri)centromere and have been assigned some roles, the understanding of their organization, their evolutionary dynamics and the disclosure of the molecular mechanisms that drive their spread, movement and rearrangement in euchromatin is still very limited. The key requirement that would help to understand the evolution and function of these sequences is their detection at the genome level and genome-wide studies of satDNAs in various model organisms. One of the main reasons for the current lack of global and in-depth studies of satDNAs in euchromatin is certainly the fact that satDNA regions are the most difficult part of the genome to sequence and assemble, and therefore they are underrepresented or absent even in the best genome assemblies (reviewed in [13,14]). However, with the improvement of long-read sequencing technology, especially Oxford Nanopore platform, it has recently been demonstrated that genomic loci made of satDNAs can be efficiently assembled [15]. Moreover, end-to-end maps of all human chromosomes, spanning highly repetitive (peri)centromeric and telomeric regions, have recently been obtained [16].

The beetle *T. castaneum* is a worldwide pest of stored products and the representative of the most species-rich animal order on Earth, the coleopterans. *T. castaneum* has become one of the most important models in the field of evolution, physiology and development of insects because its development is more representative for insects compared to *Drosophila* [17,18]. The *T. castaneum* genome has been sequenced, annotated and a first version of genome assembly was available in 2008 [19]. Structural genome analyses revealed large amounts (>30%) of different classes of repetitive DNA in the assembled part of the *T. castaneum* genome. Among them, the (peri)centromeric heterochromatic satDNA TCAST which dominates on all *T. castaneum* chromosomes comprising up to 17% of the genome [20,21], was drastically underrepresented in the assembled genome. All *T. castaneum* chromosomes are characterized by large blocks of (peri)centromeric heterochromatin, while no prominent heterochromatic blocks could be detected cytogenetically on chromosome arms [22]. Regarding the possible presence of satDNAs in euchromatin of *T. castaneum*, our previous study on the *T. castaneum* reference genome assembly disclosed that *T. castaneum* genome abounds with many different and unrelated satDNAs distributed out of (peri)centromeric regions [23]. In addition, there are also short arrays of counterpart of (peri)centromeric DNA in euchromatin regions [9]. Nine satDNAs, Cast1-Cast9, represent the most prominent non-pericentromeric satDNA portion, which was mapped to ∼0.3 % of the genome assembly at that time. In *Drosophila*, most euchromatic satDNAs feature short monomer units [24]. However, in *T. castane u*, C*m*ast1-Cast9 represent "classical" satellite DNAs with 1170 and 1340 bp long monomers. The experimentally determined genome content of Cast1-Cast9 satDNAs showed abundance of approximately ∼4.3%, which is ∼10X higher than the analyzed representativeness in the assembled genome [23]. The genome version from 2008 was based on a Sanger 7x draft assembly and was anchored to ten Linkage Groups (LGs) [19]. Recently, the genome assembly was upgraded to a new version, Tcas5.2, which was improved by using large-distance jumping libraries and BioNano Genomics optical mapping, bringing also a new official gene set OGS3 [25]. Nevertheless, our previous satDNA analyses showed that satDNA genome fractions were highly underrepresented in Tcas5.2 as well. We considered that for a deep genome-wide analysis of dominant satDNAs distributed outside the (peri)centromere, the first prerequisite would be the availability of the assembly enriched in the repetitive sequences.

In the present work, we used Oxford Nanopore Technology (ONT) long-read sequencing and applied a reference-guided assembly strategy using the Tcas5.2 reference assembly to create an improved genome assembly of *T. castaneum*. We managed to fill gaps and elongate the repetitive regions, especially the satDNAs in the new TcasONT assembly. Furthermore, we used the TcasONT assembly presented here as a platform for global and in-depth analyses of the dominant satDNA fraction in euchromatin. Our results indicate that satDNAs spread widely in euchromatic regions, including gene-rich regions. We found that these euchromatic satDNAs undergo multiple expansion events that occur on a large scale and in extensive mixed arrangements between different chromosomes. We have proposed two mechanisms for these revealed genome dynamics of these satDNAs: spread by an insertion mediated by extrachromosomal circular DNA and, to a lesser extent, spread by transposon elements (TE).

## Results

### 1. Chromosome-scale ONT-genome assembly and repetitive elements identification

The size of the reference *T. castaneum* genome assembly (Tcas5.2) is 165,9 Mb [25]. Removing placeholders and sequencing gaps results in a genome assembly size of 136 Mb. Considering that the experimentally estimated genome size is 204 Mb [26], it is clear that 68 Mb of the genome is missing from the Tcas5.2 reference assembly. In order to improve the existing assembly, especially in repetitive genome parts, we employed Oxford Nanopore Technologies’ (ONT) long-read sequencing platform. In total, 121 Gb of nanopore long reads were generated. Main steps of genome assembly process are presented in Suppl Figure 1. First, nanopore reads were divided into two fractions based on their length, short reads (<20kb; total of 52.7Gb; 258X genome coverage), and long reads (>20kb; total of 36.3 Gb; 178X genome coverage, Suppl Table 1). Next, long reads (>20kb) were used for the initial assembly using Canu [27]. This initial Canu assembly had 1479 contigs with total length of 321 Mb and a N50 of 835.5 kb (Suppl Table 2, left Canu statistics). The longest contig was 16.4 Mb in length. Our initial Canu assembly was approximately 117 Mb larger than the previously experimentally estimated genome size of 204 MB [28]. To verify the genome size and genome repetitiveness *in silico*, the k-mer frequencies of the corrected reads generated during the Canu assembly process using the FindGSE and CovEST programs were calculated. FindGSE estimated the genome to be 204 Mb, while CovEST RE showed only slightly higher estimation of 208 Mb (Supp Table 3), which is fully consistent with experimental estimation. The repeat ratio calculated by FindGSE was 27% of genome confirming the previously observed high genome repetitiveness [22]. Considering that (peri)centromeric satDNA TCAST, with 17% genome abundance, might be problematic for proper assembly [29], we assumed that the additional 117 Mb in our Canu assembly (Suppl Table 2) were primarily generated by gene-poor and highly repetitive TCAST arrays. To reduce the suspected negative effects of TCAST satDNA on the genome assembly process, we performed an additional filtering step. In this step, the contigs that did not contain at least 1000 bp of unique gene-coding sequence were removed, resulting in successful filtering of 471 out of 1479 total contigs (Suppl Figure 1).

These 471 sorted contigs were further used to achieve a new chromosome-scale assembly using reference-guided approach. The latest improved *T. castaneum* genome assembly Tcas5.2 [25] was used as the reference for organizing these contigs into chromosomes. Two reasons were crucial for the selection of this approach: 1) the availability of the high-quality *T. castane*um reference genome, Tcas5.2, and 2) assembly approach that allows the input sequence to be ordered based on the mapping quality of the reference sequence. Given that the reference genome Tcas5.2 contained 3669 unresolved gaps with total size of 11 Mb and mean gap size of 3125 Kb, the first step was included gap filling of Tcas5.2 with TGS-GapCloser using 8.6 Gb of Canu corrected larger reads (length >30 kb). TGS-GapCloser was efficient in filling most of the gaps, so in total 3607 gaps (98.3 %) were closed which increased the genome size by 10.5 Mb (Suppl table 4). The gap-filled Tcas5.2. reference genome assembly was further used as a template for orientating 471 Canu-contigs into the 10 chromosomes using RagTag software tool [30]. A total of 244 contigs were unambiguously assigned to the reference Tcas5.2. genome resulting in a new ONT-based genome assembly named TcasONT (Suppl Table 5). The rest (227 contigs) did not accurately map on the assembly, remaining assigned as unassembled sequences (Supp Table 6). To further validate structure of omitted contigs, we mapped them with highly abundant (peri)centromeric TCAST satDNAs and found that around half of unplaced sequences (14 Mb) are indeed mostly covered (>50 %) with TCAST satDNAs (Suppl Table 6). To gain insight into the actual representation of (peri)centromeric TCAST, we annotated the TcasONT assembly with TCAST satDNA and obtained 9446 monomers, accounting for 3.6 Mb (1.7%) of the genome.

The final polished TcasONT has a size of 225.9 Mb, with 191 Mb assembled in ten chromosomes and the remainder in unassembled contigs (Suppl Table 7). Compared to the reference Tcas5.2, the new TcasONT assembly shows a 45 Mb increase of total chromosome length (Suppl Table 7). Comparing TcasONT and Tcas5.2 chromosomes, the elongation of chromosome lengths ranges from 10.6 to 56.3 %, indicating a significant improvement of genome assembly continuity at the chromosome level (Suppl Table 8).

The result of completeness and quality of the new assembly revealed high levels of macrosynteny and collinearity across all chromosomes with high sequence identity between TcasONT assembly and the reference Tcas5.2 (Figure 1A). Furthermore, we calculated that 88% of the total Tcas5.2 unplaced contig sequence length is now placed within chromosomes of the new TcasONT assembly.

**Figure 1.**
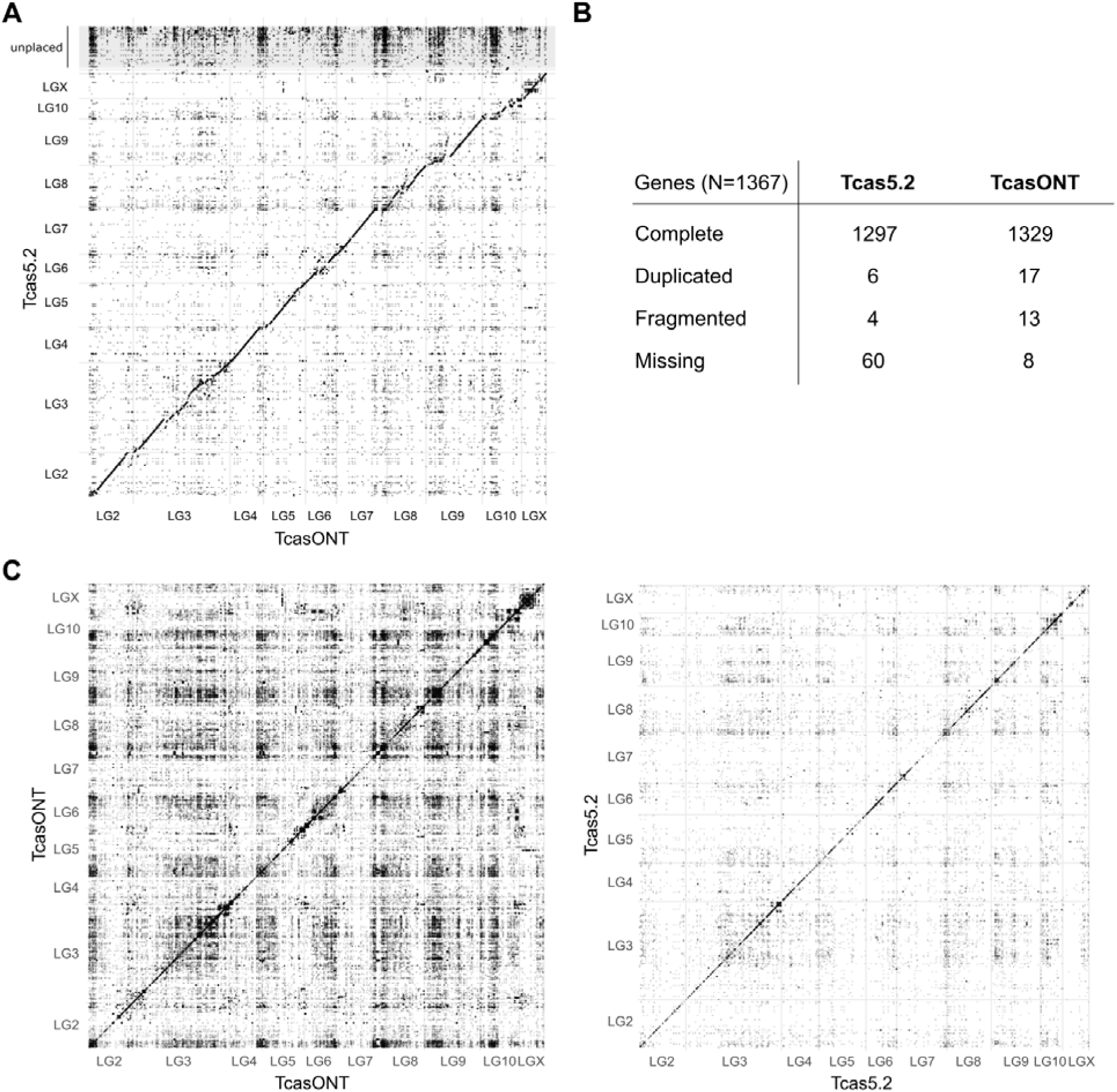
Assessment of *T. castaneum* assemblies. **A** Dot-plot representation of TcasONT vs. Tcas5.2 assembly comparison. The horizontal axis corresponds to the intervals along the TcasONT assembly, and the vertical axis corresponds to the intervals along the Tcas5.2 assembly. LG2-LG10, including LGX represent chromosomes, while unplaced contigs contain sequences in Tcas5.2. that were not associated with any chromosome. Dots closest to the diagonal line indicate co-linearity between the two assemblies. **B** Gene completeness assessment based on BUSCO analysis of Tcas5.2 and TcasONT assemblies using insect universal orthologs. The analysis was expressed as absolute numbers for complete and single-copy, complete and duplicated, fragmented and missing genes. **C** The whole genome-to-genome dot-plots analyses of TcasONT (left), and Tcas5.2 (right) assemblies. Each dot represents a region of ≥1000 bp long region which is mapped in its entirety to another part of the genome. The dot density is proportional to the number of highly similar regions.

We next assessed the gene completeness of the TcasONT assembly by comparative BUSCO analyses between TcasONT and Tcas5.2 using insect universal orthologs from the odb10 database (Figure 1B). Of the 1367 genes, we identified 1329, which corresponds to 97.2 % of BUSCO single-copy completeness. Further, 17 genes were duplicated, 13 were fragmented and only 8 genes were missing. A considerable increase in number of complete BUSCOs (32 genes) can be observed in the TcasONT assembly compared to Tcas5.2. In addition to BUSCO analysis, we “lifted” genes from the Tcas5.2 annotations onto the TcasONT assembly using the Liftoff package and assessed their final numbers (Suppl Table 9). Only 48 out of 14467 genes (0.3%) were left unmapped, mostly representing genes with no known biological functions.

To evaluate the improvement in the representation of the repetitive genome portion in the new TcasONT assembly compared to Tcas5.2, a self-similarity dot-plot analyses were performed using >70% sequence identity as a criterion. The comparison of dot-plots revealed that the TcasONT assembly has a higher level of self-similarity with medium to large dark blocks representing regions of repetitive sequences in comparison to Tcas5.2 assembly (Figure 1C). Quantified, the number of self-similar sequences was 44,671 in the TcasONT genome assembly, while in Tcas5.2 it was only 2,202, which means that the new TcasONT assembly has about 20-fold higher amount of repetitive genome fraction. To define the improvement specifically in transposable elements (TEs) content, the TEs in both Tcas5.2 and TcasONT assemblies were annotated using RepBase database and RepeatMasker. The TEs were classified into 4 main types, including DNA elements, long interspersed nuclear elements (LINEs), long terminal repeats (LTRs) and short interspersed nuclear elements (SINEs). The TcasONT assembly revealed an increase of 11,297 DNA TEs, 12,268 LTR TEs and 27,553 LINE elements (Figure 2A and Suppl Table 10) compared to Tcas5.2. Next, we investigated abundance of these elements in both genomes. Among the four TE categories, in terms of cumulative sequence occupancy there is a notable increase in TcasONT compared to Tcas5.2 for three of them; DNA TEs (5.2 Mb), LINE (14.5 Mb) and LTR (1.8 Mb) (Figure 2B and Suppl Table 10). Taken together, compared to Tcas5.2, the TcasONT assembly is enriched with 21.5 Mb of TEs (Suppl Table 10).

**Figure 2.**
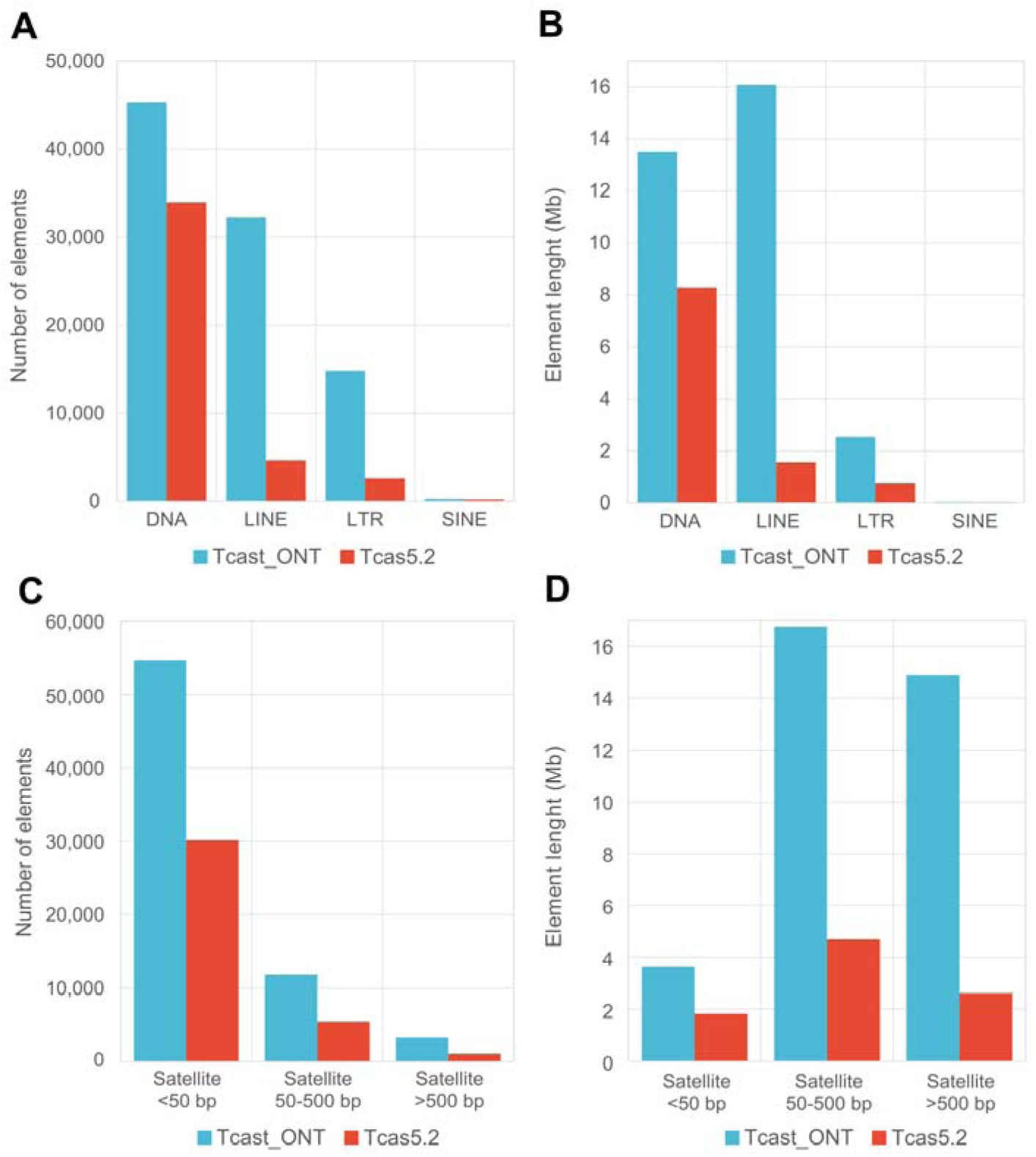
Comparison of main classes of transposable elements (TEs) found by RepeatMasker and tandem repeats (TRs) found by TRF in the TcasONT and Tcas5.2 assemblies. The height of the bars indicates **A** total number of TEs, **B** cumulative length of TEs**, C** total number of TRs, and **D** cumulative length of TRs.

Additionally, we assessed the content of another class of repetitive DNA sequences, tandem repeats (TRs), detecting them by the Tandem Repeat Finder (TRF), which was successfully used in previous studies of *T. castaneum* [23]. Detected TRs were classified into 3 groups based on their monomer length (<50bp, 50-500bp, and >500bp). The analysis revealed a total of 35.3 Mb TRs in the TcasONT assembly compared to 9.1 Mb in Tcas5.2 (Suppl Table 10), indicating a significant improvement in TRs content. On closer inspection, the results show that the number of TRs elements detected in the TcasONT assembly was doubled in all three classes (Figure 2C), while the 50-500 bp and >500 bp TRs, which represent “classical” satDNAs, contributed significantly to the cumulative genome length (31.7 Mb in TcasONT vs 7.3Mb in Tcas5.2) (Figure 2D).

From all above, the 45 Mb size difference between the Tcas5.2 and TcasONT genome assemblies is mainly due to the enrichment of repetitive regions (45.8 Mb; Suppl Table 10). The repetitive fractions that were most enriched in the TcasONT assembly were TEs (21.5 Mb) and TRs (26.2 Mb). If we consider that the (peri)centromeric TCAST satDNA accounts for only 3.6 Mb, it is evident that the TcasONT assembly has an increase of 22.6 Mb in TRs outside of (peri)centromeric regions which mainly includes an increase in “classical” satDNAs with >50 bp long monomer units.

### 2. Characterization of Cast1-Cast9 satDNAs in TcasONT assembly

Given the demonstrated improvement in the genome assembly, especially in its repetitive parts, the new TcasONT assembly represents an excellent platform for in-deep analyses of previously disclosed abundant satDNA sequences outside the (peri)centromeres, Cast1-Cast9 [23].

The first step, essential for downstream analyses, was the mapping of the Cast1-Cast9 arrays in the TcasONT assembly. Because of the well-known variability of monomer sequences within a satDNA family, it was important to establish sequence similarity and sequence coverage parameters, which would ensure the detection of the vast majority of Cast1-Cast9 monomers in the TcasONT assembly. For this reason, the BLAST search of the raw Nanopore data was performed with the consensus sequences of the Cast1-Cast9 monomers using two parameters, sequence coverage and similarity. The results were shown as density plots where color intensity correlated with number of monomers of a particular Cast satDNA (Suppl Figure 2). The majority of Cast satDNAs show the highest intensity, i.e. monomer aggregation in the area corresponding to >75% of sequence coverage and >70% of percent identity. However, the pattern with two areas of high density observed in Cast2 and Cast4 is a consequence of the mutual similarity in one part of sequences between these satDNAs (Suppl Figure 3). Despite their partial similarity they were still considered as individual satDNAs. Considering density plots for all Cast satDNAs, the parameters sequence coverage > 70% and percent identity > 70% were used to ensure mapping of most monomers of all 9 satDNAs (Cast1-Cast9).

To disclose the organization of the Cast1-Cast9 arrays, we analyzed the relationship between monomer distance in the arrays for each Cast (Suppl Figure 4). More specifically, we wanted to determine whether the arrays were organized in tandem continuously (consisting exclusively of monomers of a particular Cast satDNA) or had mixed tandem organization (with a different sequence incorporated into the arrays). The results showed that arrays of most satDNAs show no steep increases in the curve, pattern representing a typical satDNA array organization consisting of continuously arranged monomeric variants. In contrast, Cast2, Cast5 and Cast7 exhibited a disturbed tandem contiguity demonstrated by sharp increases in average array length at certain distances. The sharpest increase is visible in Cast2, indicating the presence of a specific sequence within the arrays that required further investigation. In addition to monomeric Cast2 arrays, we found Cast2 monomers also organized as a part of a new, longer repeat unit of approximately 1270 bp in length (Suppl Figure 5A-B). This new family of tandem repeats is termed Cast2’, and in the downstream analysis these two types of Cast2 arrays were examined separately. A more detailed introspection of Cast5 arrays showed that the observed profile is the result of occurrence of previously described R66-like sequences [23] dispersed within continuous Cast5 arrays (Suppl Figure 5C). In addition, analyses of Cast7 arrays also reveal a mixed organization where Cast7 monomers often occur in association with (peri)centromeric TCAST satDNA but with low sequence length and similarity (Suppl Figure 4 and 5D).

Considering the difficulty of correctly assembling satDNA sequences, the next question was whether the array lengths and arrangement of the Cast1-Cast9 satDNAs in the TcasONT assembly correspond to the real situation of the genomic loci containing the Cast1-Cast9 repeats. Namely, repetitiveness of satDNAs can lead to assembly collapse, which can result in the number of monomer units (array length) in a genome assembly being lower than in the real genome. Given that raw reads provide a realistic picture of what is originally present in the genome as they do not undergo an assembly process, we performed a comparative analysis of Cast1-Cast9 across the individual reads and across the new genome assembly. For this analysis, it was crucial to define optimal parameters for array detection to avoid fragmentation due to short gaps. Therefore, considering monomer length and mixed array organization of specific Cast satDNAs such as Cast2, Cast5 and Cast7, the best window length that ensures detection of the maximum number of arrays was evaluated for each Cast satDNA. The analysis showed that a minimum of three consecutive monomer units should be a criterion for correct array characterization for each Cast satDNA. The array patterns between the datasets of individual reads and the TcasONT assembly show remarkable similarity for almost all Cast satDNAs (Suppl Figure 6). The only exception is Cast6, whose pattern indicates a suppression of the representation of arrays in the assembly regardless of size.

To quantify the enhanced detection of Cast1-Cast9 regions in TcasONT genome assembly, we compared both their content and array length with Tcas5.2 (Table 2). In terms of overall enrichment in Cast satDNA portion, the TcasONT assembly shows an increase of 7.2 Mb in the representation of analyzed Cast satDNAs compared with Tcas5.2. Among them, Cast1, Cast2’ and Cast5 emerged as particularly prominent satDNAs with an increase of 0.7 Mb, 2.8 Mb and 2.6 Mb, respectively. The total content of Cast satDNAs in the new TcasONT genome was 8.8 Mb (4.6% of the genome), which is remarkably close to the previous experimental estimate of the genome abundance for these satDNAs (>4.3% in [23]). Among them, Cast2’ and Cast5 showed the highest genome abundance with 1.5% for each, Cast8 and Cast9 the lowest (<0.1%) while the rest remained in the moderate genome abundance level (0.1-0.4%). The results also show an exceptional increase in the number of arrays in the TcasONT assembly for Cast2’, Cast5 and Cast7. In Cast2’, for example, the number of arrays has increased by more than 9X. In addition to the number of arrays, average increase in array length (expressed as mean values) can also be observed in the TcasONT genome assembly for all Cast satDNAs. A significant increase in average array length of up to 3X was observed for Cast2’, Cast5 and Cast6 satDNAs compared to Tcas5.2. The length of an average array for Cast satDNAs in the TcasONT assembly ranges from 1.2 to 31 kb. The highest average array size of ∼31 kb has been found for Cast6, while Cast1 has the longest array of ∼112 kb.

**Table 1.** Statistics of Cast1-9 arrays detected in the Tcas5.2 and TcasONT genome assemblies.

|  | Tcas5.2 |  |  |  |  | TcasONT |  |  |  |  | Total length difference (bp) |
| --- | --- | --- | --- | --- | --- | --- | --- | --- | --- | --- | --- |
|  | Array number | Total length (bp) | Array mean (bp) | Maximal array length (bp) | genome abundance (%) | Array number | Total length (bp) | Array mean (bp) | Maximal array length (bp) | genome abundance (%) |  |
| **Cast1** | 71 | 206,931 | 2,915 | 61,324 | 0.140 | 106 | 865,816 | 8,168 | 112,523 | 0.452 | 658,885 |
| **Cast2** | 121 | 178,456 | 1,475 | 17,682 | 0.121 | 77 | 267,482 | 3,474 | 54,320 | 0.140 | 89,026 |
| **Cast2'** | 29 | 117,847 | 4,064 | 21,938 | 0.080 | 258 | 2,928,418 | 11,350 | 66,157 | 1.529 | 2,810,571 |
| **Cast3** | 173 | 220,599 | 1,275 | 4,173 | 0.149 | 172 | 246,560 | 1,433 | 10,679 | 0.129 | 25,961 |
| **Cast4** | 100 | 147,755 | 1,478 | 9,103 | 0.100 | 88 | 334,031 | 3,796 | 56,181 | 0.174 | 186,276 |
| **Cast5** | 66 | 258,820 | 3,922 | 12,025 | 0.175 | 148 | 2,897,522 | 19,578 | 85,403 | 1.512 | 2,638,702 |
| **Cast6** | 17 | 166,879 | 9,816 | 43,796 | 0.113 | 23 | 715,260 | 31,098 | 77,335 | 0.373 | 548,381 |
| **Cast7** | 86 | 82,188 | 956 | 3,729 | 0.056 | 273 | 332,266 | 1,217 | 6,028 | 0.173 | 250,078 |
| **Cast8** | 32 | 38,553 | 1,205 | 4,280 | 0.026 | 24 | 51,621 | 2,151 | 6,463 | 0.027 | 13,068 |
| **Cast9** | 117 | 145,832 | 1,246 | 4,575 | 0.099 | 98 | 139,574 | 1,424 | 5,958 | 0.073 | -6,258 |
| **Total** | 812 | 1,563,860 |  |  | 1.059 | 1267 | 8,778,550 |  |  | 4.582 | 7,214,690 |

The graph of Cast1-Cast9 monomer copy number in TcasONT compared to Tcas5.2 also clearly shows the presence of significant enrichment (2-10X) in the majority of satDNAs analyzed (Figure 3A). From the above, the new TcasONT assembly contains the representative sample of Cast arrays and can be taken as a credible platform for further genome-wide analysis of Cast satDNAs.

**Figure 3.**
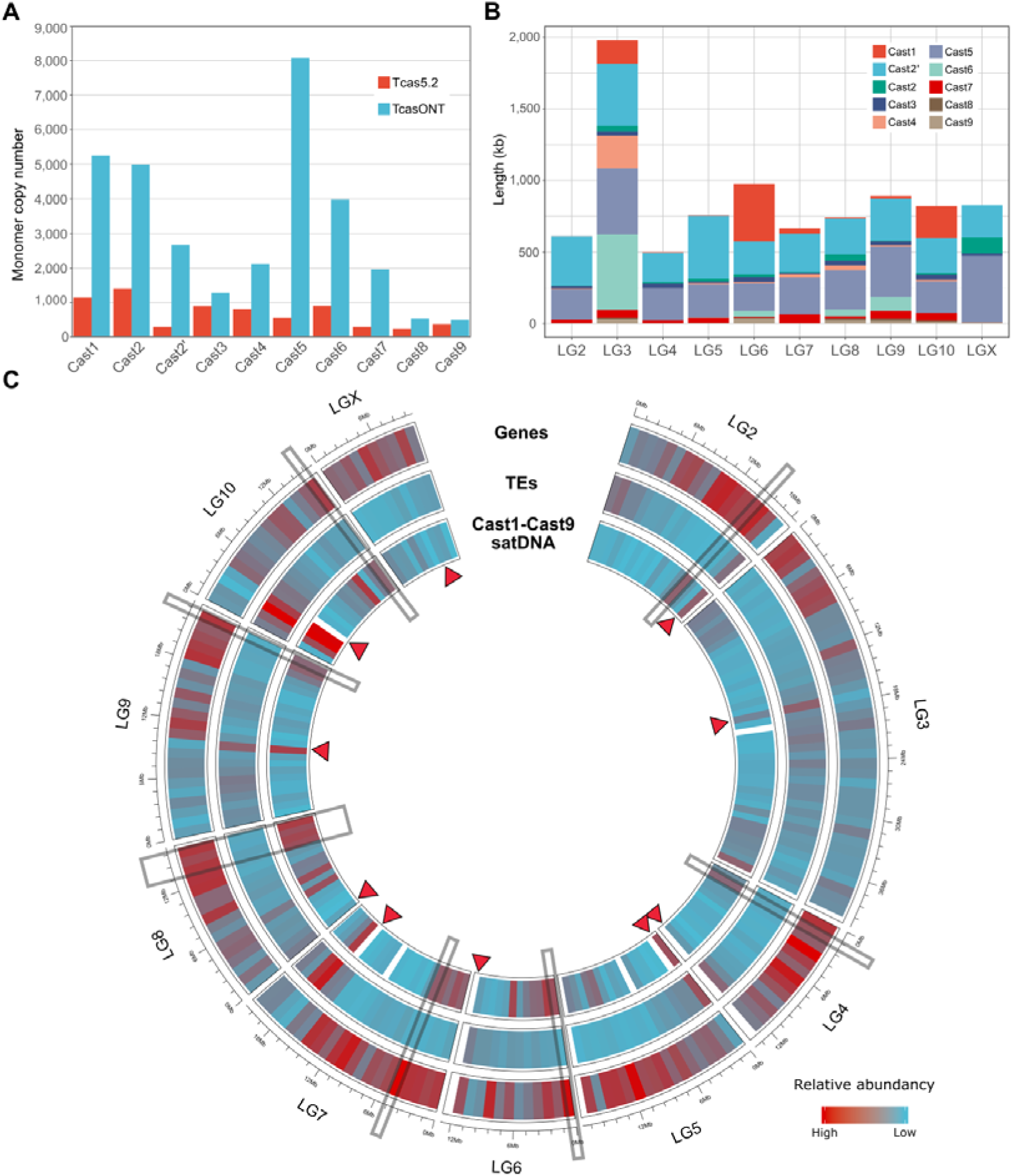
Distribution evaluation of Cast1-Cast9 satDNA in TcasONT assembly. **A** Comparison of monomer copy numbers for Cast satDNAs annotated in the reference Tcas5.2 and the TcasONT genome assemblies (see also Suppl Table 11). The assemblies were mapped with Cast1-Cast9 satDNA monomeri consensus sequences published previously [23]. **B** Comparison of monomer copy numbers for Cast satDNAs annotated in the reference Tcas5.2 and the TcasONT genome assemblies (see also Suppl Table 11). The assemblies were mapped with Cast1-Cast9 satDNA monomeric consensus sequences published previously [23]. C Circular representation of the TcasONT genome assembly with annotated Cast1-Cast9 satDNAs, genes and transposable elements (TEs). The boxes indicate the most prominent regions where the accumulation of Cast satDNAs correlates positively with gene density and negatively with the TEs density. The position of the (peri)centromere on each chromosome is indicated by a red arrow.

### 3. Cast1-Cast9 satDNAs chromosome distribution and genomic environment

The distribution of Cast1-Cast9 including Cast2’ satDNAs on *T. castaneum* chromosomes was further investigated by mapping all Cast1-Cast9 monomers on the assembled TcasONT linkage groups. The two main patterns were observed. The first pattern is characterized by Cast satDNAs located on a subset of chromosomes (Figure 3B). For example, Cast1 dominates on LG3, LG6 and LG10, while it is underrepresented on other chromosomes. Similarly, Cast6 dominates on LG3, but is also present, albeit to a much lesser extent, on LG6, LG8 and LG9. In contrast, the other Cast satDNAs such as Cast2’, Cast5 and Cast7 show a more uniform pattern characterized by similar content on almost all chromosomes (Figure 3B).

Next, we analyzed the distribution of Cast1-Cast9 along chromosomal length (Suppl Figure 7). Since it is already known that TCAST builds up huge blocks of (peri)centromeric heterochromatin [20], the presumed localization of the (peri)centromere on each chromosome is marked. Detailed examination of Cast1-Cast9 distribution disclosed an exceptional spread of Cast satDNAs along the chromosomes with no tendency to cluster in proximal regions of the (peri)centromere. Only Cast7 occurs preferentially in regions enclosed by (peri)centromeric satDNA TCAST. In addition, some Cast satDNAs show a tendency to cluster in distal chromosomal regions.

The distribution of Cast1-Cast9 satDNAs on chromosomes compared with genes and transposon elements (TEs) was also analyzed (Figure 3C). Interestingly, Cast1-Cast9 satDNAs are often part of gene-rich regions and rarely overlap with TEs (the most prominent regions with these features are highlighted in Figure 3C). The only coincidence of the occurrence of Cast satDNA and TEs was found in the (peri)centromeric regions, which corresponds to the occurrence of Cast7 satDNA.

For a more detailed analysis of Cast1-Cast9 satDNAs arrays, 50 kb flanking regions surrounding Cast1-Cast9 arrays were examined for the presence of genes and TEs. More specifically, all genes in the flanking regions of Cast1-Cast9 arrays were counted, and the base gene content of the whole genome assembly was calculated and used to assess the relative gene density in the vicinity of Cast arrays (Figure 4A). From the scaled values based on the total number of arrays, it is evident that Cast satDNAs have median gene content values that are above the median of the genome, except for Cast1, Cast6 and Cast7. Moreover, the arrays Cast2’, Cast3, Cast5 and Cast8 are surrounded by a significantly larger number of genes (Kolmogorov-Smirnov test, p<0.01) than the rest of the genome (Suppl Table 12). In addition, these Cast satDNAs have distributions above 3^rd^ quartile range of the genome indicating the presence of arrays embedded in extremely gene-rich regions. The only Cast satDNA that has a lower number of adjacent genes is Cast7 (Supple Table 12, Figure 4A). This correlates with the observation that Cast7 arrays occur in an intermingled organization with (peri)centromeric TCAST satDNA (Suppl Figure 7). In contrast, analyses of Cast satDNAs distribution in relation to TEs showed a significantly lower number of TEs near Cast satDNA arrays compared to the whole genome (Kolmogorov-Smirnov p<0.01) (Figure 4B).

**Figure 4.**
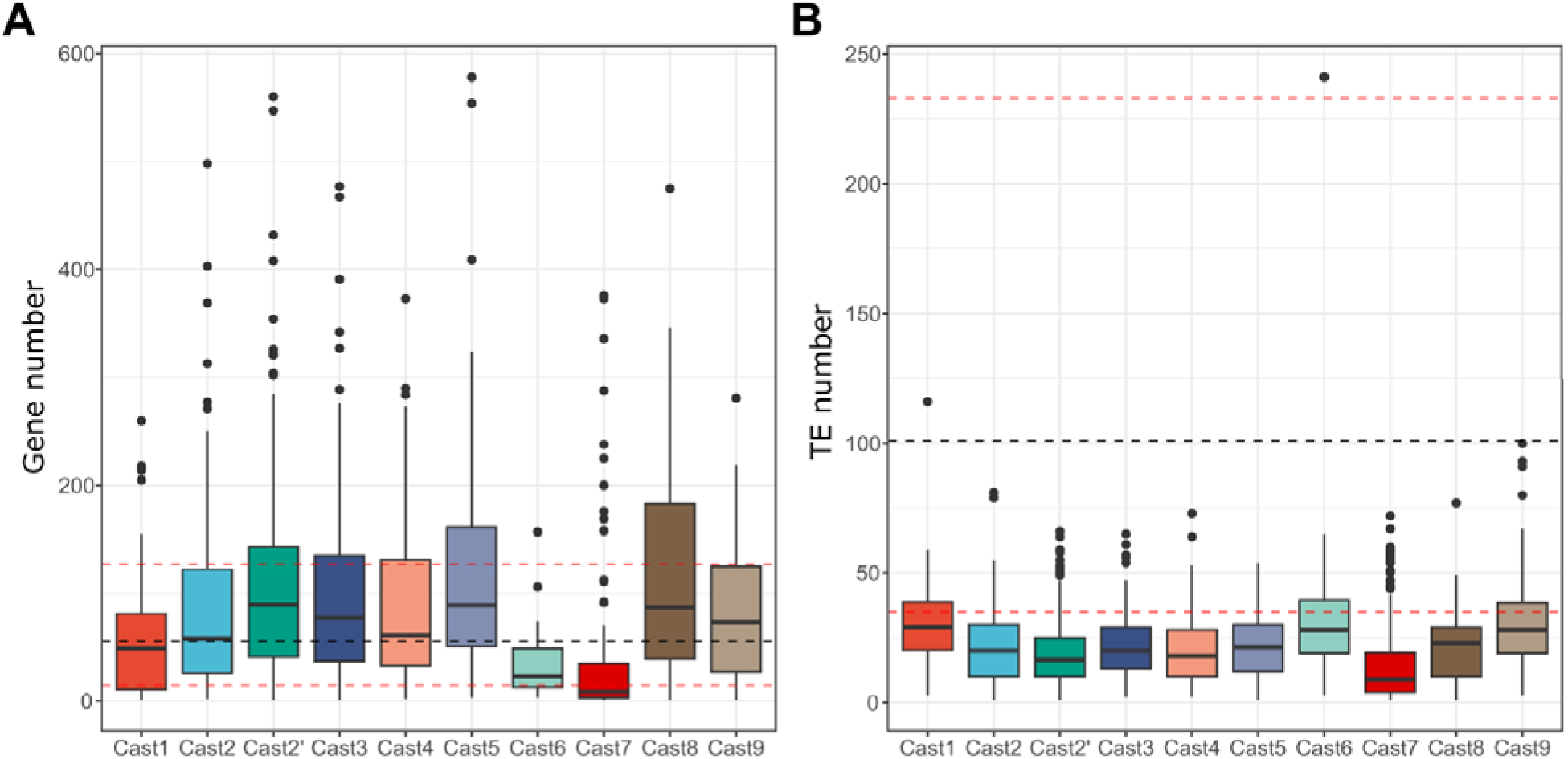
Distribution of gene and transposable element counts in vicinity of Cast arrays. **A** The number of genes was calculated based on the distribution of gene content in the assembled genome with the black line representing the median number of genes in regions of same size (100 kb windows, 5kb sliding frame) and the red dotted lines represent the first and third quartiles (Q1 and Q3) of the genome wide distribution. Results of the Kolmogorov-Smirnov tests are represented in Suppl Table 12. **B** The number of transposable elements was calculated using the same window size as for the genes, with black and red lines representing the same values. All Cast arrays had significantly (Kolmogorov-Smirnov p<0.01) less transposable elements in their vicinity when compared to the genome wide distribution, highlighting the fact that they do not overlap with TE presence.

Next, we aimed to determine if there is a correlation between the arrays’ length and the density of nearby genes. For this purpose, Cast1-Cast9 arrays were divided into three classes according to their length: (small arrays (<1000kb), moderate arrays (1000-10 000 kb) and long arrays (>10 000kb) and gene density was calculated. The results for each Cast satDNAs are presented in Suppl Figure 8. Interestingly, most Cast satDNAs have the same profile with the highest gene density being associated with arrays of intermediate length (1000-10 000 bp). In addition, a significant number of genes were observed in the Cast5 arrays, especially near large arrays (>10 kb). In the examined 50kb windows around satDNA monomers, gene counts show similar trends (Suppl Figure 8), without distinct graph patterns, meaning that the arrays are uniformly embedded in gene-rich regions.

### 4. Genomic dynamics and mechanisms of propagation of Cast satDNAs

We considered that analyzes based on the comparison of the mutual variability of monomers and arrays, together with their position on the chromosomes, can be informative for monitoring the genomic dynamics of Cast satDNAs. Taking into account that the data set (monomers and arrays) was large and the overall variability of monomers within the family relatively low (Suppl Figure 2), we noted that phylogenetic analyzes of a huge number of similar sequences would not be sufficiently sensitive to disclose trends in Cast satDNAs genome dynamics. Therefore, we decided to perform the principal component analysis (PCA) of monomer alignments, and sequence similarity relationships between the arrays for each of Cast satDNAs (Figure 5A and C). First, a genome-wide database of Cast monomers was created and annotated with their actual chromosomal positions. PCA results obtained this way showed a similar trend for almost all Cast satDNAs characterized by a scattered pattern of monomers. This is particularly evident for Cast1, Cast2, Cast2’, Cast3 and Cast4 satDNAs, where regions of extremely high density of monomers from different chromosomes become visible. The only exception with a pattern characterized by an accumulation of monomers from the same chromosome was observed in Cast6, satDNAs with long homogenized arrays.

**Figure 5.**
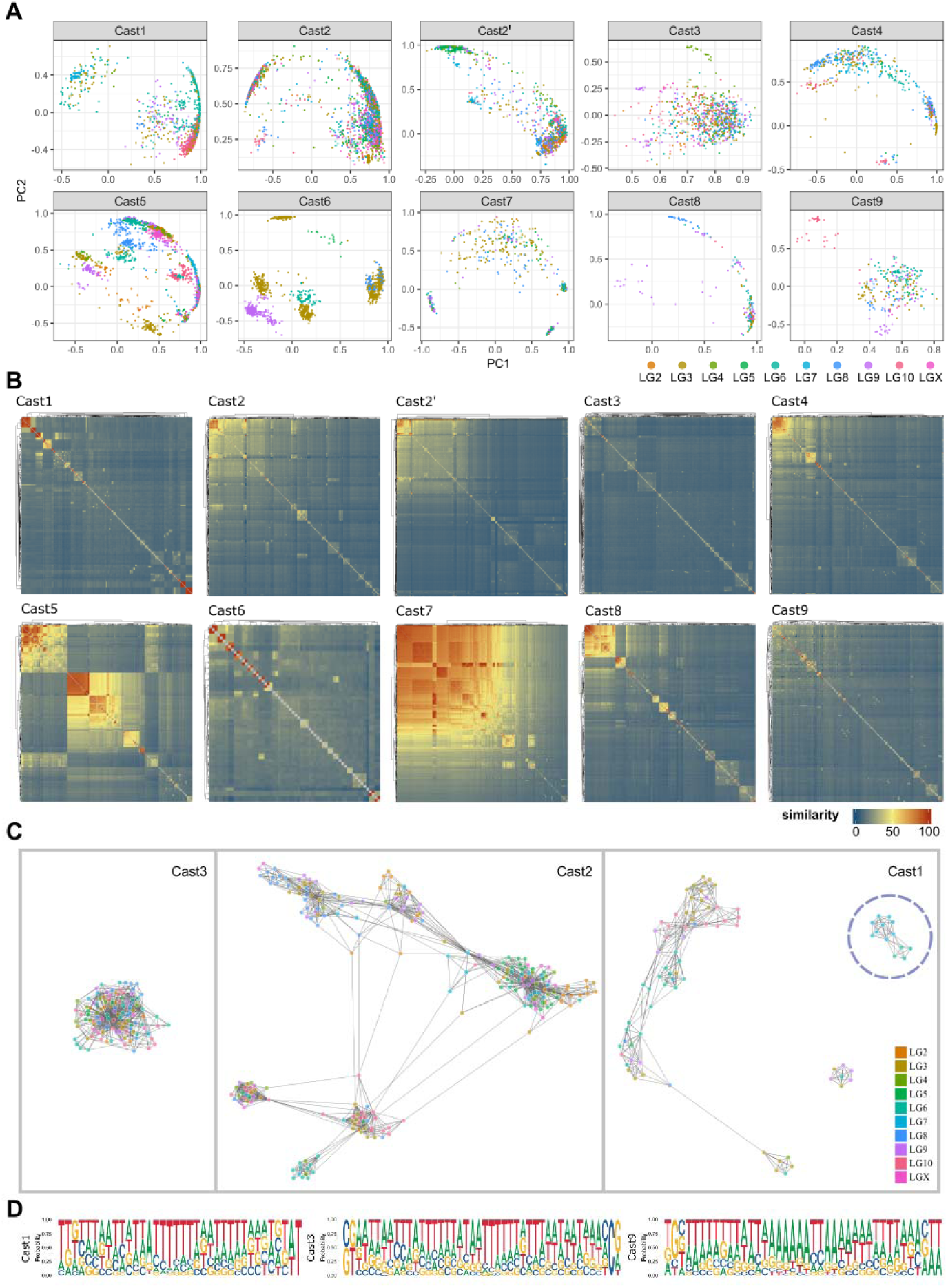
**A** Principal component analysis (PCA) of aligned Cast1–Cast9 satDNAs monomer distance matrices. Dots represent a monomer unit. Monomers were colored according to the chromosome of origin. The most prominent and recent exchange events between different chromosomes are visible in regions with high density of points with different colors. **B** Heatmap visualization of sequence similarity of 2 kb regions around Cast1-Cast9 arrays throughout the TcasONT assembly. The legend indicates that the sequence similarity of the surrounding regions increases, with colors going from blue to red. **C** Graph networks of Cast3, Cast2 and Cast1 arrays based on their sequence similarity relationship. Cast 3 shows a cluster containing all arrays. Cast2 shows a pattern with six different clusters containing arrays mixed from different chromosomes. Cast1 shows no significant clustering of arrays, except small divergent clusters. **D** Sequence logo of junction regions of Cast1, Cast3 and Cast9 arrays in which an enrichment of polyT and polyA tracts can be recognized.

Second, after a database of Cast arrays with corresponding chromosome annotation having been created, comparative analyzes of the arrays were performed based on their mutual sequence variability for each Cast satDNAs. Sequence similarity relationships between the arrays for each of Cast satDNAs are represented by graph networks (Figure 5C, Suppl Data), except for Cast6 and Cast7 which were not analyzed due to the small number of arrays. The distance and interconnectivity between dots correlate with the sequence similarity between arrays, with closer dots indicating higher similarity of monomers between different arrays. Our assumption is that the relationship between the arrays reflects their spread throughout genome. Three graphs, Cast1, Cast2, and Cast3 were selected as illustrative examples (Figure 5C), while the other Cast satDNAs arrays networks are presented in Suppl Data. The Cast3 shows the pattern of array dynamics with a dominant cluster containing relatively closely related arrays from all or almost all chromosomes, and the similar pattern applies to Cast4, Cast8 and Cast9. In addition to the dominant mixed clusters, Cast4, Cast8 and Cast9 also have some small distant clusters (up to 10 arrays) from the same chromosome. The second pattern includes Cast2 with six relatively distant array clusters with related arrays from different chromosomes, indicating multiple genome events of intense array expansion during Cast2 evolution. Interestingly, only one of them represents intrachromosomal expansion, while the rest show very extensive interchromosomal exchange. A similar pattern with several distant clusters containing related sequences from different chromosomes is also observed in Cast2’ and Cast5. Finally, the Cast1 network shows clusters with higher distance between arrays, with some array sets completely separated due to their sequence divergence. The Cast1 network also shows two separate subgroups of sequences representing different variants, one of them (Figure 5C, labelled) directly connected to the transposon element, Polytron (Suppl Figure 9).

Interestingly, these three different patterns of Cast satDNA propagation events correlate with the average lengths of the arrays. For example, satDNAs for which only one expansion event can be observed (Cast3, Cast4, Cast8 and Cast9) have a relatively short array length (mostly around 4000 bp). SatDNAs with several expansion events, as seen in Cast2, Cast2’ and Cast5, have a moderate array length of about 15000 bp. Finally, Cast1, for which no recent expansion centers were observed, intends to have very long arrays (up to 112kb).

To gain insight into the spread of Cast1-Cast9 arrays throughout the genome, the surrounding regions of the Cast arrays were also analyzed. To address this, regions of 2 kb in length upstream and downstream of individual Cast arrays were extracted. The extracted regions were aligned using the MAFFT program and the sequence similarity values were calculated from the alignment for each Cast satDNA. The results showed that of the ten Cast satDNAs analyzed, only Cast5 and Cast7 show similarity of surrounding regions for most arrays (Figure 5B). A detailed analysis of the surrounding regions for Cast5 arrays revealed two dominant regions consisting of sequences with high similarity. One of them is the R66-like sequence, found as a scattered sequence in the Cast5 arrays (see Suppl Figure 5C) being also located at one end. On the other side of most Cast5 arrays the R140-like sequence was found. Detailed analysis showed that out of 150 Cast5 arrays, 2/3 arrays had R140-like sequences and 1/3 arrays had R66-like sequences at the ends of the arrays. Likewise, Cast7 arrays showed that most of them are surrounded by the (peri)centromeric TCAST (Suppl Figure 5D). The rest of Cast satDNAs showed only partial similarities of the surrounding regions noticed only for a smaller number of arrays. (for example, Figure 5B, Cast1 panel). For example, a high-similarity subset of Cast1 arrays showed that those nine arrays originate from the same chromosome (LG7), having the same transposon-like sequence at the end of arrays (Suppl Figure 9).

For proper examination of the precise Cast array junction regions, their edges need to be precisely defined. Since the monomers at the ends of the arrays have a higher variability due to the less efficient recombination at the ends of arrays (reviewed in [14]), we developed a k-mer similarity-based approach. The results of the analyzes did not reveal any part of the monomer that would be favored at the ends of the Cast arrays (data not shown). However, the analyzes of 20 bp sequence motifs near the array boundaries showed that Cast1, Cast3, and Cast9 have regions in the vicinity of arrays with significant poly A/poly T tracts, whereas the other Cast satDNAs do not share any common motif (Figure 5D, Suppl Figure 10).

Considering the observed extensive inter- and intrachromosomal exchanges of all Cast satDNAs, one of the possible mechanisms that influence this phenomenon could be the insertion of satDNA arrays mediated by extrachromosomal circular DNA (eccDNA). To test the presence of Cast satDNAs in the eccDNA fraction, we performed two-dimensional (2D) agarose gel electrophoresis followed by Southern blot hybridization. The hybridization probes were developed for the most abundant Cast satDNAs, Cast1, Cast2’, Cast5 and Cast6 (0.3-1.5% of the genome), and the eccDNA molecules containing the examined satDNAs were indeed detected (Figure 6).

**Figure 6.**
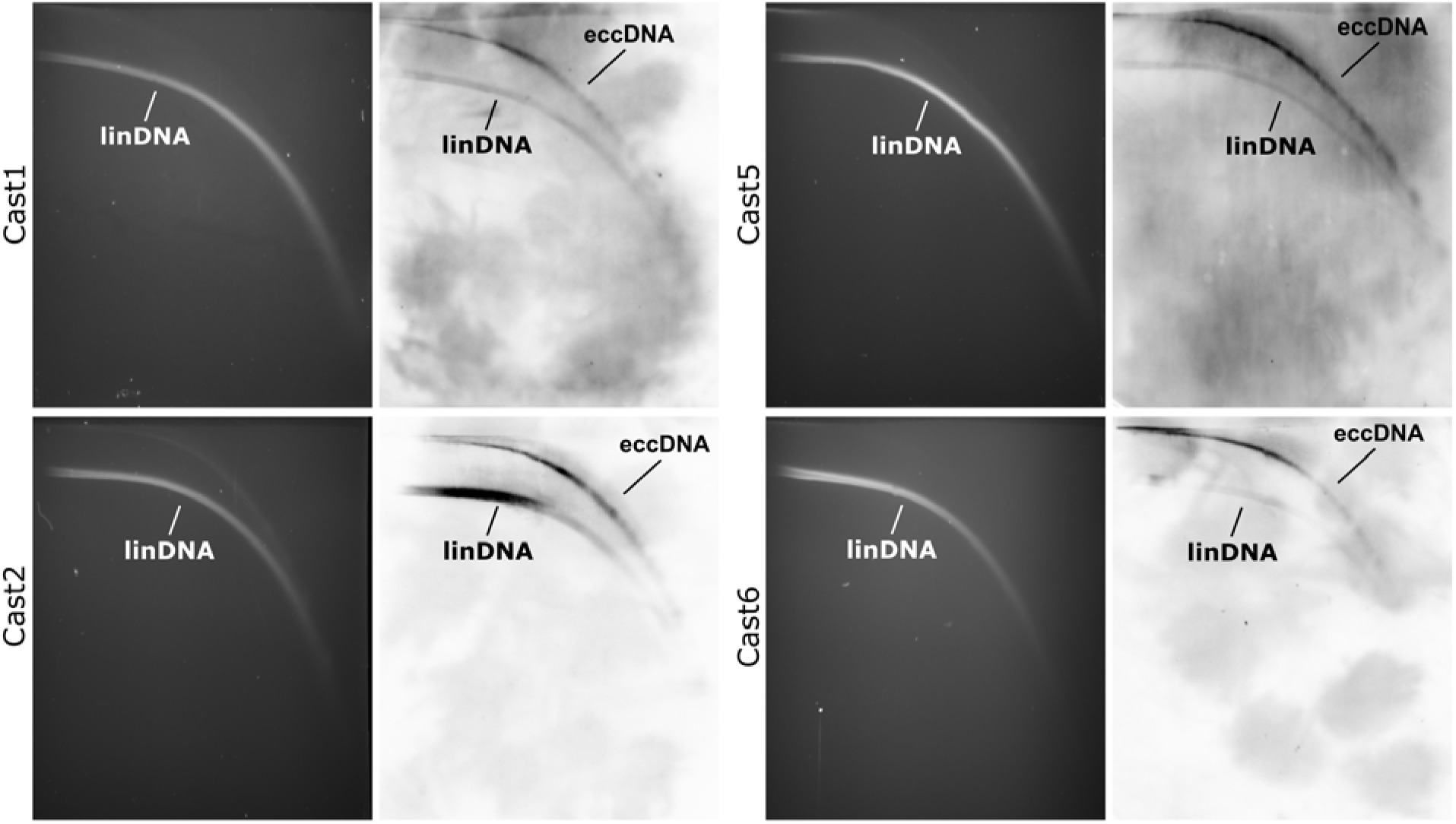
Detection of extrachromosomal circular DNA molecules containing *T. castaneum* Cast1, Cast2, Cast5 and Cast6 satDNA. Results of *T. castaneum* genomic DNA separation in 2D electrophoresis are shown on the left, while Southern blot hybridization with specific probes for each Cast element is shown on the right. Signals obtained by Southern blots confirm the presence of specific Cast satDNA in chromosomal linear double-stranded DNA (linDNA), but also in extrachromosomal circular DNA (eccDNA) molecules.

In general, chromosomal regions with suppressed or without recombination tend to accumulate satDNAs [31]. Since sex chromosomes represent regions with suppressed recombination, we examined the number and length of Cast1-Cast9 arrays on the X chromosome to compared it with autosomal chromosomes. This type of analysis would be even more informative using the Y chromosome, which is mostly non-recombining, but unfortunately, the Y chromosome was not available in the former Tcas5.2 nor in the new TcasONT assembly. Mapping of Cast1-Cast9 arrays to chromosomes disclosed that the X chromosome did not have a significantly higher average number of Cast1-Cast9 arrays per Mb, compared to the others (Supp Table 13). The comparison of the sequence variability of Cast satDNA monomers from X chromosome in contrast to autosomes did not show significant difference. However, the analysis of Cast1-Cast9 array length showed a statistically significant longer length of arrays on the X chromosome compared to the others (Wilcoxon test, p<0.05) (Figure 7). This trend is especially prominent in Cast 2, Cast2’, Cast5 and Cast9, where the average length of the arrays is increased even 10X at the X chromosome (e.g. Cast2’) (Suppl Figure 11).

**Figure 7.**
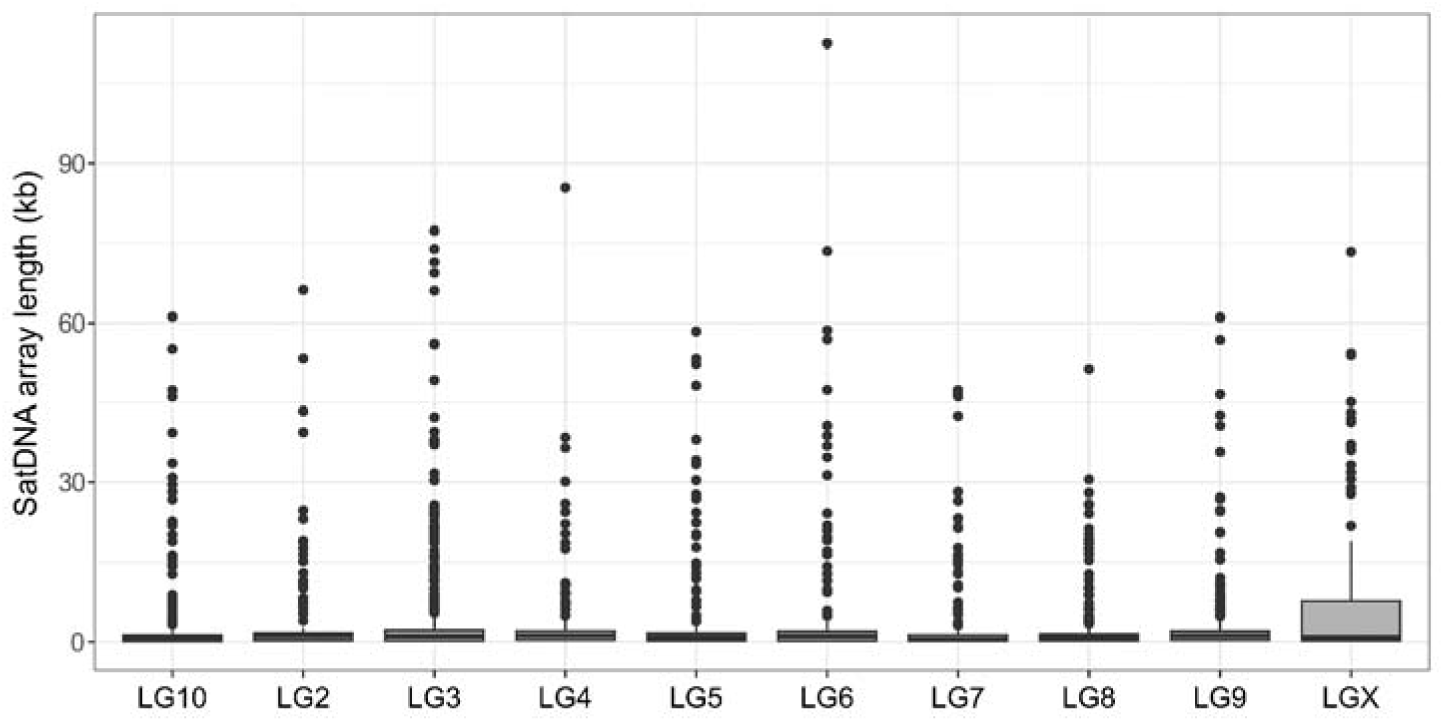
Distribution of Cast1-Cast9 satDNA array lengths across chromosomes, LG2-LG10 including X chromosome (LGX).

## Discussion

Despite significant advances in sequencing technologies and plethora bioinformatics tools, satDNAs are still the most difficult part of a genome to assemble. The most illustrative example of the problem with satDNA in assembly process using long-read sequencing is found in the recently published work on holocentric nematodes [32]. Although these genomes are small, the assemblies were quite fragmented, and the authors pointed out that abundant and scattered satDNAs occupying holocentromeres are the main problem preventing assembly at the chromosome level [32,33]. It is clear that the assembly of satDNAs-rich genomes is challenging, even when long-read sequencing is used. Previous studies [22,23] indicated that *T. castaneum* is one of the species with a considerable amount of diverse repetitive sequences, and therefore it is not surprising that the official reference assembly, although of good quality in the coding part, was significantly deficient in long-range assembly of repetitive DNAs.

The aim of our research was a thorough investigation of satDNAs in *T. castaneum* euchromatin, and for this purpose we needed an assembly of reliable quality in repetitive regions. Unfortunately, the official assembly Tcas5.2 did not fulfill this criterion, and we decided to improve the *T. castaneum* assembly, focusing on upgrading the difficult-to-assemble repetitive DNA regions. To this goal, a high-quality *T. castaneum* genome assembly at the chromosome level was generated by combining nanopore long-read sequencing and a reference-guided approach. The new TcasONT assembly is nearly 45 Mb larger than the latest version of the Tcas5.2 reference genome. The achieved better completeness of genes compared to the reference Tcas5.2 underlines the credibility of TcasONT for gene representation. The TcasONT assembled chromosomes lack only 13 Mb of the estimated *T. castaneum* genome sequence of 204 Mb, previously determined experimentally [28] and also *in silico* in our study. The missing 13 Mb could primarily be attributed to (peri)centromeric regions, due to assembly-impeding highly repetitive TCAST satDNA regions [21]. Despite significant advances in sequencing technologies and assembly methods, it is not unusual that (peri)centromeric regions remain still incomplete even in most nanopore-based assemblies [34]. The only exceptions are the recently published Telomere-to-Telomere assemblies of prominent model organisms such as human and *Arabidopsis* that contain the complete (peri)centromere [16,35,36]

In addition to enrichment in the gene regions, TcasONT assembly proved to be highly representative for the study of the repetitive regions. Namely, comparison of the assemblies showed a 20-fold enrichment in the repetitive part in the TcasONT assembly compared to Tcas5.2. In TcasONT, the representation of TEs has also been increased, both in terms of the number of elements and the abundance. However, the increase is particularly significant for tandem repeats (TRs) with a monomer unit length of >50 bp, which are declared as “classical” satDNA. The enrichment of the proportion of TEs and TRs totals 47.8 Mb compared to Tcas5.2. Our study proves that the nanopore sequencing approach enables the resolution of previously difficult-to-assemble regions, most of which consist of highly repetitive sequences such as satDNAs and TEs. It is also important to note that TcasONT is larger than the previous version by 25%, which means that a substantial portion of the genome is now available for further research.

The assembly process often does not assemble the satDNAs correctly, resulting in collapsed arrays [29], but our comparative analyses of the raw data and the TcasONT assembly show that the TcasONT assembly is highly credible in terms of representativeness of Cast satDNAs. The TcasONT assembly showed Cast satDNAs abundance of 8.8 Mb, which accounts for 4.6% of the genome, being consistent with their experimentally confirmed genome abundance [23]. The previous assembly Tcas5.2 contained only 1/10 of experimentally estimated amount of Cast satDNAs. Next, we used the TcasONT assembly provided here as a source for a comprehensive and in-depth analysis of the ten most dominant satDNAs in euchromatin, Cast1-Cas9, previously characterized in [23]. Our genomic analysis of these satDNAs included their distribution, correlations with genes and TEs, genomic dynamics, and possible mechanisms of spread in euchromatin.

Interestingly, we found that these ten "classical" satDNAs (Cast1-Cast9, including a new version of Cast2, Cast2’) do occur in the form of long tandem arrays in euchromatin, a region in which the accumulation of abundant satDNAs is not as common as in the (peri)centromeric heterochromatin. Moreover, some Cast satDNAs arrays show a very high average length (Cast6 with 31kb) and maximum length (Cast1 with 112kb). The question arises whether the presence of such long satDNA arrays in regions that are not considered suitable for the accumulation of satDNAs, is indeed a distinctive feature of the *T. castaneum* genome. We suspect that the presence of relatively numerous and abundant satDNAs in euchromatin is not a unique feature of the *T. castaneum* genome, but that the absence of satDNAs in these regions in some other genome assemblies is a consequence of significant gaps caused by assembling difficulties. We therefore believe that the strategy of nanopore sequencing, combined with reference-guided assembly approach, ensures an enviable representation of satDNAs, which should be crucial for comprehensive and in-depth satDNA studies in other species.

Furthermore, one of the important questions that arose from the observed organization, characterized by long arrays in non-pericentromeric regions, is whether Cast1-Cast9 satDNAs were truly euchromatic. Indeed, it is possible that satDNAs, even if located outside the (peri)centromeric region, could be localized in gene-poor regions populated by, for example, transposons or other types of noncoding DNA [37]. However, our gene density analyses showed that the surrounding regions of the arrays of almost all Cast satDNAs correlated positively with the gene-rich regions compared to the average gene density in the genome. Moreover, some satDNA arrays were surrounded by extremely gene-rich regions. It is also interesting that these regions are not characteristic of short arrays as expected, but long satDNA arrays (>10kb) are also found near the genes.

It has not been seen before that satDNAs in euchromatin, including those with long arrays, were mostly integrated into an environment rich in genes and poor in transposons. Contrary to expectations, we found that the occurrence of Cast1-Cast9 satDNAs and TEs often do not overlap. This fact clearly speaks of the ability of Cast satDNA to invade euchromatic regions and indicates the flexibility of euchromatin, i.e., gene-rich regions, to tolerate and accommodate these sequences. Previous models support the fact that the accumulation of satDNA is characteristic of regions where recombination is suppressed, such as (peri)centromeric heterochromatin [38]. In addition, a commonly accepted view is that genome recombination events ultimately result in array loss, and therefore satDNAs mainly persist in heterochromatic regions (reviewed in [14]). However, from our study, it appears that gene-rich euchromatin, where recombination is expected to occur, may tolerate the accumulation of satDNA. Further, comparative analysis of Cast satDNA arrays between the X and autosomal chromosomes showed that suppressed recombination in the X chromosome stimulates the propagation of satDNAs into longer arrays. However, it has no significant impact on the number of arrays and sequence variability. We propose that recombination in the euchromatin does not prevent the satDNA extensive spread but discourages arrays prolongation. This impact on arrays’ length in euchromatin is probably due to rearrangement potential of satDNAs elongation, which can affect neighboring regions. Considering that previous studies have mainly reported short satDNA arrays in euchromatin regions, our finding of "classical" satDNAs, widely present across the euchromatin, in the form of long arrays in gene-rich regions, provides new insights into the genomic distribution of satDNAs. Previous studies were based on euchromatic satDNA with short arrays, suggesting that these satDNAs may play roles in gene regulation by acting as “evolutionary tuning knobs” [35], regulating chromatin [9,12], and facilitating X chromosome recognition/dosage compensation [11]. SatDNAs located in euchromatic regions could also regulate gene expression by modulating local chromatin structure, or via transcripts derived from the repeats. For example, contractions of the human subtelomeric satellite D4Z4 alter the chromatin state of nearby genes, causing muscular dystrophy [40]. Recent studies disclosed that expression of euchromatic satDNA-derived transcripts control embryonal development in the mosquito via sequence-specific gene silencing [41]. Being in the gene-rich regions, epigenetic histone marks of satDNAs in euchromatin could also influence surrounding gene regions. For example, human genome-wide analysis of satDNAs in euchromatin has shown their association with repressive histone mark H3K9me3, suggesting that they may affect the expression of neighboring genes [42]. In addition to the direct impact on genes, large-scale rearrangements that involve the long Cast1-Cast9 arrays scattered throughout the genome, are also highly probable. The ability of euchromatic satDNA to undergo high evolutionary turnover may also contribute to the rapid change of gene landscape across the genome, and consequently on gene function [43]. In summary, the presence of such a substantial portion of Cast1-Cast9 satDNAs in euchromatin regions of the *T. castaneum* genome can strongly influence both gene expression and the dynamics of genome evolution.

Another important emerging issue, given the widespread distribution of Cast1-Cast9 satDNAs in euchromatin and their putative impact on the genome evolution, is the way in which these satDNAs are propagated. While the mechanisms of TE propagation are well known and depend on the type of transposon elements [44], the propagation of satDNA, especially in euchromatin regions, is still quite a mystery. Given that we know little of the genome dynamics and molecular mechanisms that shape the distribution of euchromatic satDNA on the genome scale, we carried out analysis of the distribution pattern, genome dynamics as well as analysis of surrounding and junction regions of Cast1-Cast9 satDNAs. The results mainly show arrays dispersed on all chromosomes and along their entire chromosome length without regional preference. Interestingly, the results of the sequence similarity relationships between arrays analyzed at the genome level for each Cast satDNAs, showed grouping of Cast satDNAs in three characteristic patterns (shown in Figure 5C). If we assume that the observed patterns of satDNA behavior are events captured at a specific and distinct point of “satDNA life” in the genome, here we propose a course of events that can explain the genome dynamics of euchromatic satDNAs over time. The timeline of possible events is presented in a simplified form, as shown in Figure 8. The first event could be extensive expansion from one center, when short arrays dramatically spread and shuffled among different chromosomes (t1). After expansion, the arrays of the satDNAs diverged in sequence due to their different localization and reduced effect of homogenization. At the same time, if arrays are located in a stimulating environment for the propagation, their arrays become lengthened (t2). After this phase, some of the arrays, probably short ones, could be roused and form new expansion centers, characterized again by a rapid spread to different chromosomes (t3). After many expansion events, satDNA could enter a dormant state in terms of spreading but continue to extend arrays (t4). In a favorable environment, the arrays can be subjected to extensive elongation and homogenization. It is difficult to determine what causes a particular sequence to become a promoter of dispersal. However, the observed pattern of euchromatic satDNA genome dynamics suggests an efficient mechanism of their extensive and efficient spread on different chromosomes.

**Figure 8.**
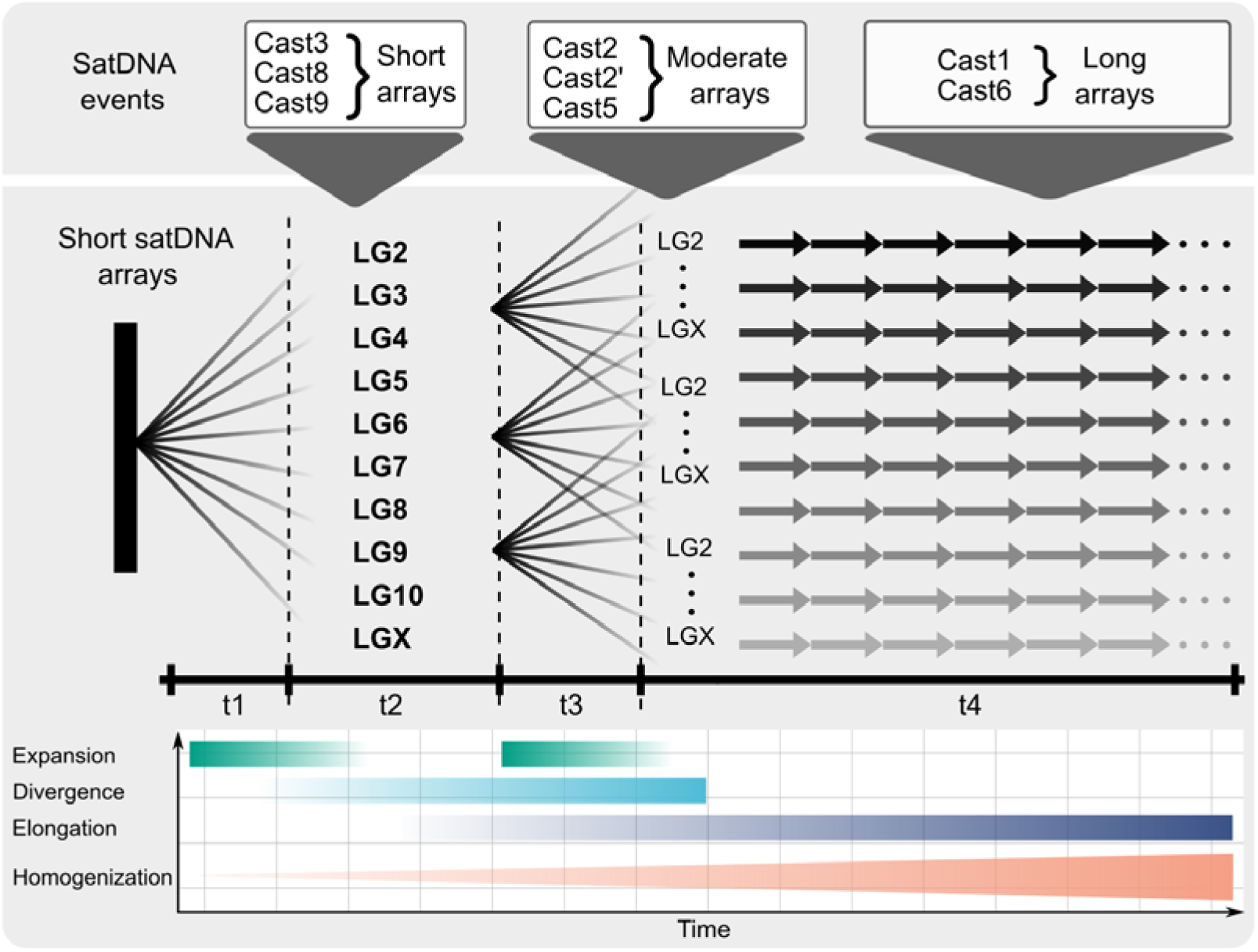
The most parsimonious scenario of genome dynamics of euchromatic Cast satDNAs. The multiple expanding lines extending from a single genomic locus represent a sudden burst event in which satDNA spreads on different chromosomes, followed by a second expansion event that leads to further propagation of satDNA. Evidence of such behavior in *T. castaneum* is shown on top of the figures with three different examples of euchromatic satDNA. The timeline depicts the main events in different colors. Expansion is described as multiple sudden bursts denoted by a sudden increase in intensity and subsequently followed by a slow decrease. The divergence event lasts longer, and its effect is more prominent with time. Elongation has an even later onset with a similarly increasing effect over time. Homogenization is a permanent process affecting all satDNAs, however the impact of homogenization is the strongest on the longest arrays.

Although satDNAs were considered less mobile compared to TEs, which move extensively throughout the genome ultimately forming interspersed repeats, our study showed that satDNAs also have very high capacity to spread throughout the genome. It is common for TEs to spread through the genome in recurrent bursts, which is characterized by the movement of many TEs through the genome within a short evolutionary time [45]. Our results show that euchromatin satDNAs exhibit a similar pattern of genomic dynamics, which includes the cycles of repeated bursts. Although it is assumed that spreading bursts of TEs can occur for no apparent reason, it has been shown that they are often associated with stressful situations such as extreme temperatures, radiation, chemical exposure, or viral infections [45]. In a view of a similar expansion pattern with TEs, it can be speculated that euchromatic satDNAs can also enter in expansion cycles triggered by an external cause such as stressful living conditions. Another indication that stress can affect satDNAs in euchromatin of *T. castaneum* is the finding that euchromatin counterparts of pericentromeric TCAST satDNAs are more strongly expressed as response to the heat stress [12].

To date, the three mechanisms have been proposed to be involved in the propagation of satDNA through the euchromatin: (i) dispersion in a form of short arrays integrated as central repeats of non-autonomous transposons elements [46]; (ii) the spread of satDNAs across long physical distances in euchromatin through eccDNAs [47]; (iii) interlocus gene conversion events via 3D interactions between loci in interphase nucleus [48].

There are a few examples where dispersed satDNAs have been found in the form of short arrays integrated as central repeats of non-autonomous transposons elements [46,49,50]. Moreover, tandem repeats within Tetris transposon DNA have been reported to be the basis for generating long satellite arrays in *Drosophila virilis,* that eventually lose transposon features in adjacent regions [46]. Considering that most of the Cast5 arrays have transposon-like sequences in adjacent regions, one of which shows low similarities with the Mariner element, it is highly probable that Cast5 satDNA owes its expansion to this element. However, other Cast satDNAs, except in some subsets, did not show a tendency to associate with other mobile elements, but show very extensive propagation, indicating an efficient self-propagation mechanism that operates at the inter- and intra-chromosome level. Mechanisms that have recently been highlighted in *Drosophila* as possible drivers of satDNAs spread throughout euchromatin in the X chromosome, are eccDNA reintegration and interlocus gene conversion [43]. To disclose the putative mechanisms involved in the spread of euchromatin of satDNA, we analyzed junction regions. The ends of the arrays, which could represent a critical sequence for the insertion of the array into the environment, did not show a regular pattern, suggesting that gene conversion does not represent mechanisms primarily involved in the spread. We found, however, that some Cast satDNAs have a preferential insertion site in regions characterized by the presence of poly-A/T tracts with microhomology. On the other hand, Cast satDNAs were shown to be present in the eccDNA fraction, suggesting that the spread of Cast satDNA may be mediated by eccDNA. Based on the analyses of neighboring regions, junctions, eccDNA fraction and considering the observed extensive spread of satDNA, we proposed the two mechanisms responsible for euchromatin satDNAs genome dynamics and evolution: transposition and eccDNA insertion.

In conclusion, this work has achieved two important accomplishments: (1) a high-quality reference genome TcasONT of the insect model organism *T. castaneum*, 2) an in-deep genome-wide analysis of the dominant euchromatic satDNAs using the new *T. castaneum* assembly. First, by using Oxford Nanopore long-read sequencing, we have produced the most contiguous genome assembly of *T. castaneum* to date, with significant improvement in the representation of the repetitive genome portion and making a significant step toward completing the genome assembly of this scientifically and economically important model organism. Our reference genome assembly has a great potential to be important genomic resources for future research of both genes and various repetitive fractions of the genome. Second, using the new assembly, our comprehensive analyses of the satDNAs, embedded in euchromatin regions, revealed their massive spread and proliferation in gene-rich regions, challenging the current hypothesis of a precluded coexistence of genes and abundant satellite DNA. Based on the analyzed dynamics of satDNA spreading through the euchromatin, we suggested the evolutionary scenario characterized by the recurrent burst, implying movement on the different chromosomes within a short evolutionary time. Then, we propose two mechanisms that are most likely involved in efficient propagation of satDNAs in euchromatin: transition by TEs and/or insertion by eccDNA. In summary, such dynamical sequences embedded in euchromatin, which are subject to changes and rearrangements, would have an extraordinary potential for rapid evolution of the genome and consequently of the species itself. This opens a new perspective on satDNAs by considering them as inevitable parts of euchromatin, thus stimulating new research involving transcriptomic and epigenetic studies, which could disclose their role and putative influence on gene content.

## Materials and methods

### Insects sample and DNA preparation

Laboratory cultures of the red flour beetle *T. castaneum*, highly cultured strain Georgia 2 (GA2), were routinely reared in whole wheat flour with the addition of whole rye flour and oats under conditions favoring faster multiplication (32 °C and 70% relative humidity) in the dark. Selection of the different life stages of the insects was performed by sieving through a 0.71-mm sieve and picking the beetles individually with tweezers. Larvae and pupae were collected to achieve the most efficient DNA isolation efficiency. The collected amount of beetles had to be sufficient to perform the isolation with 200 mg of the starting material for pupae and 500 mg for larvae as starting sample for each DNA isolation. DNA isolation was performed following the protocol described in [46]. Briefly, to isolate DNA of sufficient quality and quantity, nuclei were first isolated using a nuclei isolation buffer (NIB), then lysis, and finally transferred to a Qiagen Genomic Tip 100/G column. Pure DNA was spooled from eluates and precipitated in isopropanol. In total, we have performed 8 of the described isolations, 4 for larvae and 4 for pupae.

### Library preparation and sequencing

Oxford Nanopore library was prepared using SQK-LSK110 library preparation kit. Preparation was performed according to the manufacturer’s instructions with modifications described below. Prior to sequencing DNA was sheared 30X times using a 30-gauge needle and purified of short fragments using the Short Read Eliminator (SRE) XS kit (Circulomics) according to the manufacturers protocol. The sheared and purified DNA was then used in library preparation protocols in amounts 2X larger than the ONT protocol recommendation. Additionally, all recommended elution and incubation times were extended to twice the length. These steps were performed to avoid library loss and to increase final library concentration sufficient to load multiple MinION flow cell (final concentrations of the library were >100 ng/µL allowing multiple loads from a single library). The flow cells were washed and reloaded 2-5 times to maximize the output. A total of 7 MinION 10.3.4 and 9.4.1 flow cells were used to obtain the data with a total cumulative output of 89.9 GB and an N50 of 20.1 kb. The software used for sequencing was the Oxford Nanopore MinKnow 20.10.3.

### Genome assembly

Data obtained from all sequencing runs was basecalled using Oxford Nanopore basecalling software Guppy 5.0.1 with parameters as listed in the Suppl Table 14. The basecalled reads were then used in the assembly process, using Canu v2.2 [26] with the parameters listed in the Suppl Table 14. According to Canu documentation the parameters were adjusted for high repetitiveness of the genome [22] and the high observed AT content in the reference Tas5.2 genome assembly [24]. The reads were filtered to >20 kb to reduce the computational intensity needed to perform the assembly, given the amount of data and the relatively small genome of the beetle. Incorporating only those reads that had a mean quality score of >7 and a length >20kb, a total of 44 Gb was used for the new genome assembly representing genome coverage of 224X (Suppl Table 1). Selected reads were up to 500 kb in length with a mean read length of 47 kb and mean quality score per read of 12.8. Canu assembly was performed using the resources of computer cluster Isabella based in University Computing Centre (SRCE), University of Zagreb.

### Filling the gaps

To successfully arrange the Canu contigs into chromosomes based on the Tcas5.2 (GCF_000002335.3) assembly, pre-existing gaps in the Tcas5.2 assembly needed to be bridged. This was done using the TGS-Gapcloser software [47] with default for gap-filling settings (Suppl Table 14) using corrected reads generated as the output of the Canu pipeline. Gap filling was used to bridge small and medium gaps in the Tcas5.2 assembly to prevent Nanopore-based long contigs from being interrupted due to discontinuity of the previous assembly.

### Contig orientation

After filling the gaps in the Tcas5.2 assembly chromosomes, the RagTag collection of software tools [29] was used. The contigs obtained with Canu were used as the query sequence and the gap-filled Tcas5.2 assembly was used as the reference sequence with “scaffold” parameter (Suppl Table 14). RagTag uses the reference sequence (gap filled Tcas5.2 assembly), as a guide to which it aligns and places the input sequences (Canu contigs) to create the new assembly and fill the gaps in the subject assembly with new successfully aligned contigs. Since the highly repetitive regions are in majority lacking from the Tcas5.2 assembly, this approach allowed to incorporate a new region of repetitive elements in the TcasONT assembly. RagTag maps the high confidence genomic regions on chromosomes and then, if a contig ends or begins with a repetitive region it is placed in the gap giving us insight into previously unknown regions. The output of the RagTag tool is the unpolished assembly that is used as a template for polishing (Suppl Figure 1).

### Polishing

To improve the quality of the assembly and reduce the error ratio, correction of Canu contigs was performed. To polish the assembly, two separate rounds of RACON [48] polishing were performed using reads excluded from the initial assembly (<20 kb). These short reads used for polishing involved significant addition of genomic region information as this excluded data was ∼50 Gb. Polishing was performed according to the instructions within the RACON documentation. The RACON workflow consists of mapping the reads onto the assembled genome with minimap2 [49] and then using the information about the mapped reference reads to perform polishing. Produced assembly, named TcasONT was used for downstream analysis.

### BUSCO analysis

Benchmarking Single Copy orthologs (BUSCO) [50] analysis was performed using BUSCO v5.0.0. module on the Galaxy web platform (usegalaxy.org). Same settings, as listed in the Suppl Table 14 were repeated for all assembly validations.

### Repeat detection

RepeatMasker is a widely used tool for finding and masking different repeat elements within a given target sequence [51]. RepeatMasker was used to get the GFF/GTF formatted data with the position and orientation of classified RepBase repeat elements. This was a base for the information about the quantity, size, and distribution of different elements within examined genome assemblies. Assemblies were annotated with different elements using the RepeatMasker program on the Galaxy web platform (usegalaxy.org) with different repeat elements from the latest RepBase database (RELEASE 20181026) of repeats and the “Hexapoda” species listing for clade specific repeats. We also rerun RepeatMasker for the Tcas5.2 assembly to update the repeat annotations, as it was created using previous version of RepBase database. For the quantification of three satDNAs classes (with defined monomer length >50; 50-500; >500bp) in TcasONT and Tcas5.2 assembly, Tandem Repeat Finder (TRF) program [52] was used with default parameters.

### Gene annotation

In order to map genes in Canu contigs (filtering) and in TcasONT assembly, we used the novel *LiftOff* package [53] using gene annotations from the Tcas5.2 assembly. The Liftoff first maps the newly created TcasONT assembly to the reference Tcas5.2 and then uses these overlaps to accurately align genes from a reference Tcas5.2 assembly to a target TcasONT. This approach was limited to finding possible new genes in the improved version of the TcasONT assembly, but because the Tcas5.2 assembly was annotated based on a large database of RNA-seq reads combined with gene and general trait prediction methods, this approach ensures gene completeness.

### Identification of Cast1-Cast9 satDNAs

Annotation of satellite repeats within the genomes was performed using the standalone NCBI’s BLAST algorithm, and the interface to the R programming language package *metablastr* [54]. The subject sequences were the different analyzed assemblies (Tcas5.2 and TcasONT) while the queries were the previously characterized Cast1-Cast9 [23]. Post-analysis included filtering BLAST hits in the genome to discover the Cast1-Cast9 trends and arrays.

### Analysis of Cast1-Cast9 satDNA arrays

All Cast1-Cast9 monomers were defined from the BLAST result table, filtered according to parameters described in Suppl Figure 2. Next, it was crucial to define optimal parameters for satDNA arrays detection to avoid fragmentation due to putative short different sequence(s) incorporated into the arrays. Total arrays of particular Cast satDNA were analyzed to evaluate the best neighboring window length that ensures the connection of the continuous repeating monomers into one array (Suppl Figure 3). This approach to array definition was introduced to allow for errors and array insertions and to correctly link all existing monomers of a given satellite as accurately as possible. Afterwards we performed basic filtering to define arrays and remove short, interspersed monomers using the following parameters:

*(array!="Cast2-mix" & width>530) | (array=="Cast2-mix" & width > 2000)*

These parameters ensured arrays with 3 or more repeat units for each satDNA. The exception was the Cast2’ array (Cast2 monomer interspersed with the newly discovered sequence Cast2’) where 3 different length monomers were included, out of which 1100 bp Cast2’ is mixed with 170 bp Cast2.

### Analysis of gene content in vicinity of Cast1-Cast9 arrays

A region of 50 kb upstream or downstream of the arrays was selected to define gene profiles around the different Cast1-Cast9 arrays. In these 100-kb regions, 100 equal bins of 1 kb size (50 upstream and 50 downstream of a particular array) were created, and the number of exons was counted in each of them. This allowed gene profiling around different Cast1-Cast9 arrays. Expected exon densities were calculated by computing median, 1Q, and 3Q exon densities across the entire genome in 100-kb windows using a custom R script.

### Multiple sequence alignment and PCA

We used MAFFT [56] to create multiple sequence alignments of Cast1-9 monomers in the assembly. After alignment we used the “F81” genetic distance evolutionary model from the package *ape* [57] in R to generate genetic distance matrices which were subsequently used in the PCA analysis. PCA was performed using the FactoMineR [58] package PCA function and the first 2 dimensions for each satDNA were visualized using *ggplot2*.

### Tandem array region detection

To determine exact edges of Cast1-9 arrays in the genome, we employed a specific strategy. Conventional monomer detection strategy uses a hard cutoff, based on monomer similarity, to determine their relative positions. As edges of the arrays often contain degenerate monomers, discovery of small homology and potential junction regions is often very difficult. Therefore, our strategy was based on several steps. First, a database of all monomers of particular satDNA in the genome was created together with a database of all arrays with their flanking regions (500 bp). To encompass all possible variations of satDNA sequences, all k-mers (32 bp) were extracted from monomers. Also, k-mers were extracted based on position in extracted arrays with flanks. For each position in the extended array the k-mer from the monomer database, the closest match based on Hamming distance was found and the score recorded. Next, rolling mean position score was calculated based on the mean of +/-5 position scores. Real edges were determined by finding the minimum and maximum position for each array with distance lower than 5. Based on these new edges, we extracted the surrounding and microhomology regions.

### Visualizations and calculations

All plots and calculations were produced in R using custom made data processing notebooks. Apart from the standard libraries, we also used the *circilize* package [59] to create the circular visualization plots of global genome patterns. To create the complex heatmaps used in the characterization of neighboring regions similarity we used the *ComplexHeatmap* package [60]. Utilizing a graph-based visualization method, we addressed the challenge of low variation among satDNA monomers and their tendency for intra- and interchromosomal exchange and homogenization, as illustrated by the mixing observed in the PCA plots. For each monomer in each array, we found the closest 5 monomers which do not belong to the same array using *dist.dna* function from the *ape* package in R and “F81” genetic distance model. Afterwards, we produced a graph network visualization using the *networkD3* package. Clustered and connected dots in the network represent possible satDNA arrays involved in frequent exchange via some of the proposed mechanisms contrary to the disconnected nodes. Homology of 20 bp regions before and after arrays was visualized using *ggseqlogo* package in R, after alignment with MAFFT.

### Extrachromosomal circular DNA analysis on two-dimensional agarose gel electrophoresis

Two-dimensional agarose gel electrophoresis was performed according to [51], with modifications. Total DNA was isolated from 500 mg *T. castaneum* pupae by standard phenol-chloroform extraction and dissolved in Tris-EDTA buffer, pH=8.0. The concentration was measured with the Qubit 4 fluorometer (Invitrogen). 20 µg of DNA was passed 25 times through a 0.33 mm hypodermic needle to shear the linear DNA. Given that the proportion of linear double-stranded DNA (dsDNA) fragments in the total genomic DNA (gDNA) isolate greatly exceeds the amount of potential extrachromosomal DNA (eccDNA) molecules, we treated the total gDNA isolate with exonuclease V, which cleaves linear dsDNA in both directions starting at both 5’ and 3’ termini of linear dsDNA. The sheared DNA was digested with exonuclease V (New England Biolabs) overnight at 37 °C to remove as much as possible linear dsDNA fragments and keep the circular DNA intact. The reaction was stopped by adding 11 mM EDTA pH 8.0 and incubated at 70 °C for 30 minutes. The DNA was then purified using the Monarch PCR & DNA Cleanup Kit (NEB). The first-dimension electrophoresis was run in 0.7% agarose in 1xTBE buffer at 0.7 V/cm for 18h. After electrophoresis, the agarose gel was stained with 1xTBE buffer containing 0.2 µg/ml ethidium bromide. The lane with the separated DNA was cut out and the 1.5% agarose in 1xTBE containing 0.2 µg/ml ethidium bromide was poured around the lane, which was positioned at a 90° angle to the first electrophoresis. The second dimension was run at 4 V/cm for 3 h.

### Southern blot hybridization

To ensure effective transfer of DNA from the agarose gel to the membrane, the gel was rinsed with 0.25M HCl for 30 minutes and then with 0.4M NaOH for a further 30 minutes. The DNA was transferred to a positively charged nylon membrane (Roche Life Science) overnight by capillary transfer. Hybridization probes for the Cast1, Cast2, Cast5 and Cast6 satellite DNA were labelled by biotin-16 dUTP (Jena Biosciences) using PCR amplification of the cloned plasmids containing specific satDNA sequences. Amplification was performed with satellite-specific primers (Cast1: 5’ AAGTCGGCTACGACTAACCGTTC 3’ and 5’ TTGCAAATTTGGATTCCGCCCGG 3’; Cast2: 5’ TATACGCAAAATGAGCCGC 3’ and 5’ AAAGTCGTAGAGCAATGCGG 3’; Cast5: 5’ GGTGTTGAAAAGTCATAARTTGAGTG 3’ and 5’ AGAGCCGGTGTACACAACATT 3’; Cast6: 5’ CGACGCATGGGTCAATCTAAGACA 3’ and 5’ ATTCGAAACTTTTCAAAAAAATTGG 3’). Hybridization was performed as described by [23]. Signals were detected using streptavidin-alkaline phosphatase and the chemiluminescent substrate CDP-star (Roche Life Science) and blots were visualised using the Alliance Q9 Mini (Uvitec) imager.

## Data avaliability

The *T. castaneum* genome assembly TcasONT has been deposited to the European Nucleotide Archive (ENA) under the BioProject accession PRJEB61413 and assembly accession number GCA_950066185. The genome annotation and supplementary data presented in this study are available in FigShare at https://doi.org/10.6084/m9.figshare.22683325.

## Acknowledgments

We thank Raul Horvat for critical reading of the manuscript.

## References

1. Plohl M, Meštrović N, Mravinac B. Centromere identity from the DNA point of view. Chromosoma. 2014; doi: 10.1007/s00412-014-0462-0.

2. Blattes R, Monod C, Susbielle G, Cuvier O, Wu JH, Hsieh TS, et al.. Displacement of D1, HP1 and topoisomerase II from satellite heterochromatin by a specific polyamide. EMBO J. 2006; doi: 10.1038/sj.emboj.7601125.

3. Aldrup-MacDonald ME, Kuo ME, Sullivan LL, Chew K, Sullivan BA. Genomic variation within alpha satellite DNA influences centromere location on human chromosomes with metastable epialleles. Genome Res. 2016; doi: 10.1101/gr.206706.116.

4. Melters DP, Bradnam KR, Young HA, Telis N, May MR, Ruby JG, et al.. Comparative analysis of tandem repeats from hundreds of species reveals unique insights into centromere evolution. Genome Biol. 2013; doi: 10.1186/gb-2013-14-1-r10.

5. Bersani F, Lee E, Kharchenko P V., Xu AW, Liu M, Xega K, et al.. Pericentromeric satellite repeat expansions through RNA-derived DNA intermediates in cancer. Proc Natl Acad Sci U S A. 2015; doi: 10.1073/pnas.1518008112.

6. Ferree PM, Prasad S. How Can Satellite DNA Divergence Cause Reproductive Isolation? Let Us Count the Chromosomal Ways. Genet Res Int. 2012; doi: 10.1155/2012/430136.

7. Feliciello I, Pezer Ž, Kordiš D, Madarić BB, Ugarković D. Evolutionary history of alpha satellite DNA repeats dispersed within human genome euchromatin. Genome Biol Evol. 2020; doi: 10.1093/gbe/evaa224.

8. Kuhn GCS, Küttler H, Moreira-Filho O, Heslop-Harrison JS. The 1.688 repetitive DNA of *Drosophila*: Concerted evolution at different genomic scales and association with genes. Mol Biol Evol. 2012; doi: 10.1093/molbev/msr173.

9. Brajković J, Feliciello I, Bruvo-MadWarić B, Ugarković DW. Satellite DNA-like elements associated with genes within euchromatin of the beetle *Tribolium castaneum*. G3 Genes, Genomes, Genet. 2012; doi: 10.1534/g3.112.003467.

10. Menon DU, Coarfa C, Xiao W, Gunaratne PH, Meller VH. siRNAs from an X-linked satellite repeat promote X-chromosome recognition in *Drosophila melanogaster*. Proc Natl Acad Sci U S A. National Academy of Sciences; 2014; doi: 10.1073/pnas.1410534111.

11. Joshi SS, Meller VH. Satellite Repeats Identify X Chromatin for Dosage Compensation in *Drosophila melanogaster* Males. Curr Biol. Elsevier Ltd.; 2017; doi: 10.1016/j.cub.2017.03.078.

12. Feliciello I, Akrap I, Ugarković Đ. Satellite DNA Modulates Gene Expression in the Beetle Tribolium castaneum after Heat Stress. PLoS Genet. 2015; doi: 10.1371/journal.pgen.1005466.

13. Miga KH. Completing the human genome: the progress and challenge of satellite DNA assembly. Chromosom Res. Springer Netherlands; 2015; doi: 10.1007/s10577-015-9488-2.

14. Lower SS, McGurk MP, Clark AG, Barbash DA. Satellite DNA evolution: old ideas, new approaches. Curr Opin Genet Dev. Elsevier Ltd; 2018; doi: 10.1016/j.gde.2018.03.003.

15. Miga KH, Koren S, Rhie A, Vollger MR, Gershman A, Bzikadze A, et al.. Telomere-to-telomere assembly of a complete human X chromosome. Nature. Springer US; 2020; doi: 10.1038/s41586-020-2547-7.

16. Nurk S, Koren S, Rhie A, Rautiainen M, Bzikadze A V., Mikheenko A, et al.. The complete sequence of a human genome. Science (80- ). American Association for the Advancement of Science; 2022; doi: 10.1126/science.abj6987.

17. Klingler M, Bucher G. The red flour beetle T. castaneum: elaborate genetic toolkit and unbiased large scale RNAi screening to study insect biology and evolution. Evodevo. BioMed Central; 2022; doi: 10.1186/s13227-022-00201-9.

18. Campbell JF, Athanassiou CG, Hagstrum DW, Zhu KY. *Tribolium castaneum*: A Model Insect for Fundamental and Applied Research. https://doi.org/101146/annurev-ento-080921-075157. Annual Reviews; 2022; doi: 10.1146/ANNUREV-ENTO-080921-075157.

19. Richards S, Gibbs RA, Weinstock GM, Brown S, Denell R, Beeman RW, et al.. The genome of the model beetle and pest *Tribolium castaneum*. Nature. 2008; doi: 10.1038/nature06784.

20. Gržan T, Despot-Slade E, Meštrović N, Plohl M, Mravinac B. CenH3 distribution reveals extended centromeres in the model beetle *Tribolium castaneum*. Malik HS, editor. PLoS Genet. 2020; doi: 10.1371/journal.pgen.1009115.

21. Ugarković D, Podnar M, Plohl M. Satellite DNA of the red flour beetle *Tribolium castaneum* - Comparative study of satellites from the genus *Tribolium*. Mol Biol Evol. 1996; doi: 10.1093/oxfordjournals.molbev.a025668.

22. Wang S, Lorenzen MD, Beeman RW, Brown SJ. Analysis of repetitive DNA distribution patterns in the *Tribolium castaneum* genome. Genome Biol. 2008; doi: 10.1186/gb-2008-9-3-r61.

23. Pavlek M, Gelfand Y, Plohl M, Meštrović N. Genome-wide analysis of tandem repeats in *Tribolium castaneum* genome reveals abundant and highly dynamic tandem repeat families with satellite DNA features in euchromatic chromosomal arms. DNA Res. Oxford Academic; 2015; doi: 10.1093/dnares/dsv021.

24. Flynn JM, Long M, Wing RA, Clark AG, Arkhipova I. Evolutionary Dynamics of Abundant 7-bp Satellites in the Genome of *Drosophila virilis*. Mol Biol Evol. 2020; doi: 10.1093/molbev/msaa010.

25. Herndon N, Shelton J, Gerischer L, Ioannidis P, Ninova M, Dönitz J, et al.. Enhanced genome assembly and a new official gene set for *Tribolium castaneum*. *BMC Genomics*. BMC Genomics; 2020; doi: 10.1186/s12864-019-6394-6.

26. Brown SJ, Henry JK, Black IV WC, Denell RE. Molecular genetic manipulation of the red flour beetle: Genome organization and cloning of a ribosomal protein gene. Insect Biochem. 1990; doi: 10.1016/0020-1790(90)90011-I.

27. Koren S, Walenz BP, Berlin K, Miller JR, Bergman NH, Phillippy AM. Canu: Scalable and accurate long-read assembly via adaptive κ-mer weighting and repeat separation. Genome Res. 2017; doi: 10.1101/gr.215087.116.

28. Alvarez-Fuster A, Juan C, Petitpierre E. Genome size in *Tribolium* flour-beetles: Inter-and intraspecific variation. Genet Res. 1991; doi: 10.1017/S0016672300029542.

29. Tørresen OK, Star B, Mier P, Andrade-Navarro MA, Bateman A, Jarnot P, et al.. Tandem repeats lead to sequence assembly errors and impose multi-level challenges for genome and protein databases. Nucleic Acids Res. Oxford University Press; 2019; doi: 10.1093/nar/gkz841.

30. Alonge M, Lebeigle L, Kirsche M, Aganezov S, Wang X, Lippman ZB, et al.. Automated assembly scaffolding elevates a new tomato system for high-throughput genome editing. bioRxiv.:2021.11.18.4691352021;

31. Stephan W. Nonlinear Phenomena in the Evolution of Satellite. Berichte der Bunsengesellschaft für Phys Chemie. 1986; doi: 10.1002/bbpc.19860901119.

32. Mota APZ, Koutsovoulos GD, Perfus-Barbeoch L, Despot-Slade E, Labadie K, Aury J-M, et al.. Unzipped genome assemblies of polyploid root-knot nematodes reveal unusual and clade-specific telomeric repeats. 2024; doi: 10.1038/s41467-024-44914-y.

33. Despot-Slade E, Mravinac B, Širca S, Castagnone-Sereno P, Plohl M, Meštrović N. The Centromere Histone Is Conserved and Associated with Tandem Repeats Sharing a Conserved 19-bp Box in the Holocentromere of Meloidogyne Nematodes. Mol Biol Evol. 2021; doi: 10.1093/molbev/msaa336.

34. Talbert PB, Henikoff S. The genetics and epigenetics of satellite centromeres. Genome Res. 2022; doi: 10.1101/gr.275351.121.

35. Naish M, Alonge M, Wlodzimierz P, Tock AJ, Abramson BW, Schmücker A, et al.. The genetic and epigenetic landscape of the *Arabidopsis* centromeres. Science (80- ). 2021; doi: 10.1126/science.abi7489.

36. Altemose N, Logsdon GA, Bzikadze A V., Sidhwani P, Langley SA, Caldas G V., et al.. Complete genomic and epigenetic maps of human centromeres. Science. NIH Public Access; 2022; doi: 10.1126/SCIENCE.ABL4178.

37. Meštrović N, Mravinac B, Pavlek M, Vojvoda-Zeljko T, Šatović E, Plohl M. Structural and functional liaisons between transposable elements and satellite DNAs. Chromosom Res. 2015; doi: 10.1007/s10577-015-9483-7.

38. Charlesworth D, Charlesworth B, Marais G. Steps in the evolution of heteromorphic sex chromosomes. Heredity (Edinb). 2005; doi: 10.1038/sj.hdy.6800697.

39. King DG, Soller M, Kashi Y. Evolutionary tuning knobs. Endeavour. 1997; doi: 10.1016/S0160-9327(97)01005-3.

40. Zeng W, De Greef JC, Chen YY, Chien R, Kong X, Gregson HC, et al.. Specific loss of histone H3 lysine 9 trimethylation and HP1γ/cohesin binding at D4Z4 repeats is associated with facioscapulohumeral dystrophy (FSHD). PLoS Genet. 2009; doi: 10.1371/journal.pgen.1000559.

41. Halbach R, Miesen P, Joosten J, Taşköprü E, Rondeel I, Pennings B, et al.. A satellite repeat-derived piRNA controls embryonic development of *Aedes*. Nature. Springer US; 2020; doi: 10.1038/s41586-020-2159-2.

42. Vojvoda Zeljko T, Ugarković Đ, Pezer Ž. Differential enrichment of H3K9me3 at annotated satellite DNA repeats in human cell lines and during fetal development in mouse. Epigenetics and Chromatin. BioMed Central; 2021; doi: 10.1186/s13072-021-00423-6.

43. Sproul JS, Khost DE, Eickbush DG, Negm S, Wei X, Wong I, et al.. Dynamic evolution of euchromatic satellites on the x chromosome in *Drosophila melanogaster* and the simulans clade. Mol Biol Evol. 2020; doi: 10.1093/MOLBEV/MSAA078.

44. Kelleher ES, Barbash DA, Blumenstiel JP. Taming the Turmoil Within: New Insights on the Containment of Transposable Elements. Trends Genet. NIH Public Access; 2020; doi: 10.1016/J.TIG.2020.04.007.

45. Mérel V, Boulesteix M, Fablet M, Vieira C. Transposable Elements in *Drosophila melanogaster*. Mob DNA II. Wiley;

46. Dias GB, Svartman M, Delprat A, Ruiz A, Kuhn GCS. Tetris is a foldback transposon that provided the building blocks for an emerging satellite DNA of *Drosophila virilis*. Genome Biol Evol. 2014; doi: 10.1093/gbe/evu108.

47. Cohen S, Yacobi K, Segal D. Extrachromosomal circular DNA of tandemly repeated genomic sequences in *Drosophila*. Genome Res. 2003; doi: 10.1101/gr.907603.

48. Lee YCG, Ogiyama Y, Martins NMC, Beliveau BJ, Acevedo D, Wu C ting, et al.. Pericentromeric heterochromatin is hierarchically organized and spatially contacts H3K9me2 islands in euchromatin. PLoS Genet. 2020; doi: 10.1371/journal.pgen.1008673.

49. Vojvoda Zeljko T, Pavlek M, Meštrović N, Plohl M. Satellite DNA-like repeats are dispersed throughout the genome of the Pacific oyster *Crassostrea gigas* carried by Helentron non-autonomous mobile elements. Sci Rep. Nature Publishing Group; 2020; doi: 10.1038/s41598-020-71886-y.

50. Macas J, Neumann P. Ogre elements - A distinct group of plant Ty3/gypsy-like retrotransposons. Gene. 2007; doi: 10.1016/j.gene.2006.08.007.

51. Cohen S, Lavi S. Induction of Circles of Heterogeneous Sizes in Carcinogen-Treated Cells: Two-Dimensional Gel Analysis of Circular DNA Molecules. Mol Cell Biol. 1996; doi: 10.1128/mcb.16.5.2002.

