## Supplementary Figures for "Long-read genome assembly of the insect model organism *Tribolium castaneum* reveals spread of satellite DNA in gene-rich regions by recurrent burst events"

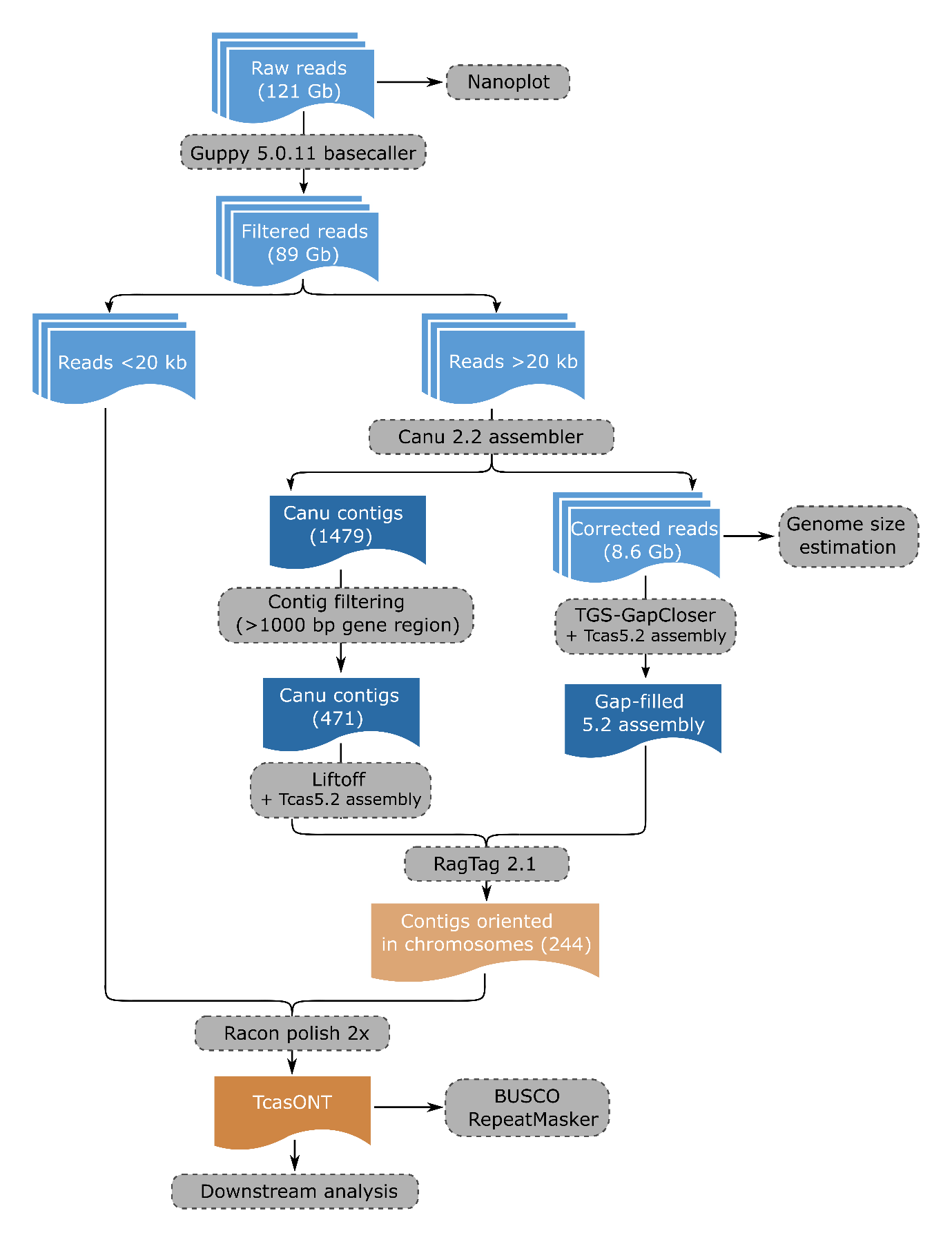

**Supplementary figure 1.** Schematics of main steps performed during genome assembly process.

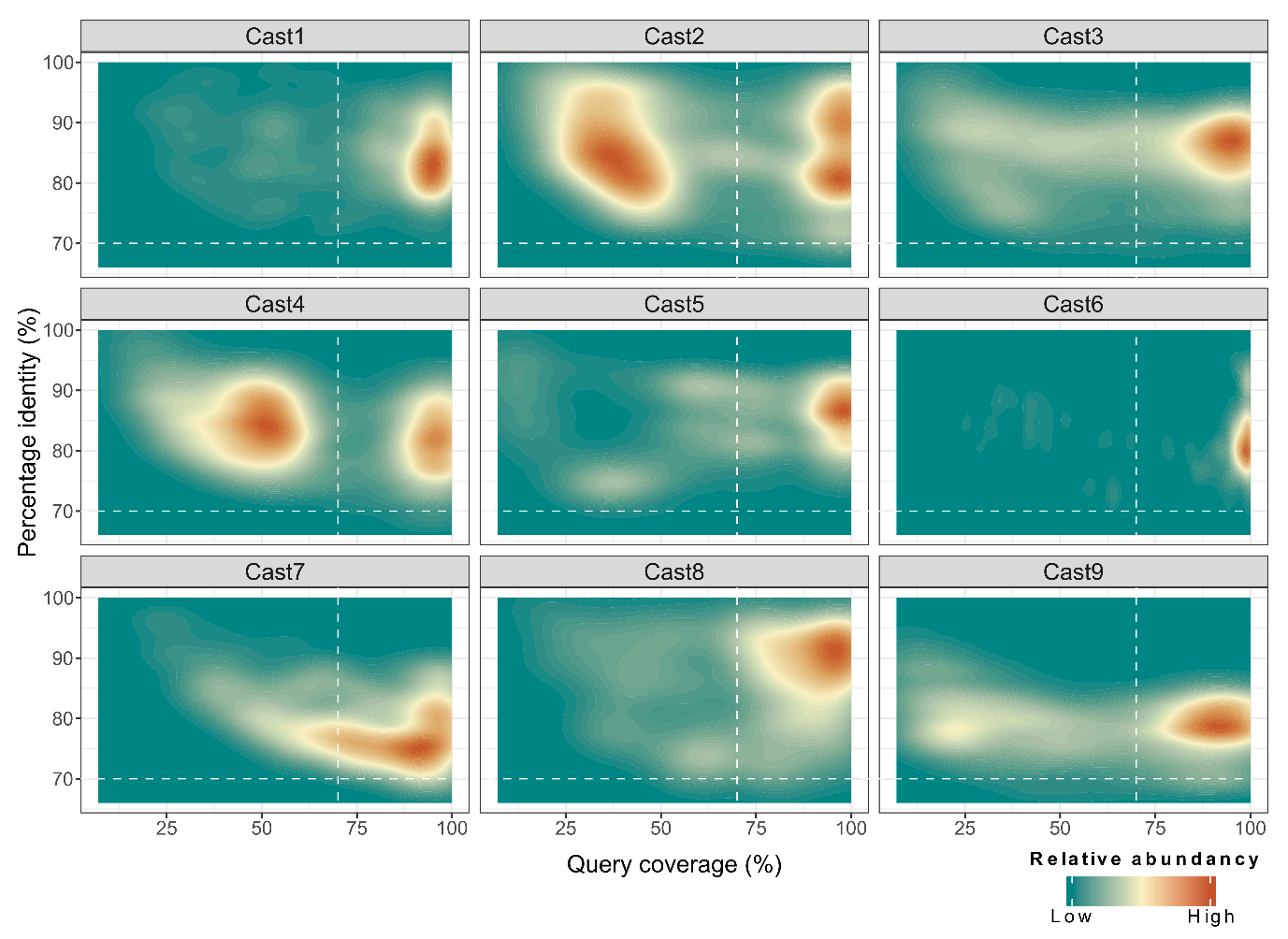

**Supplementary figure 2.** The density plot of Cast monomers of nine Cast satDNAs (Cast1-Cast2) detected by BLAST search of the TcasONT assembly. The X-axis represents the percent coverage of the query, and the Y axis represents the percent identity of the query. The relative abundance of the monomers in the graph correlates with a color gradient (from green to red). Dashed white lines represent set parameters for satDNA detection in further analysis.

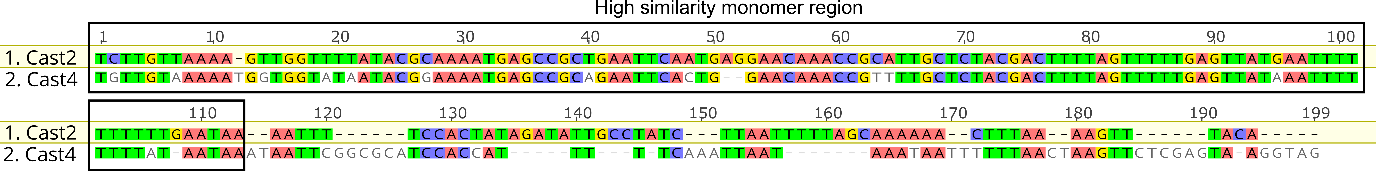

**Supplementary figure 3.** Pairwise alignment of Cast2 and Cast4 sequence with highlighted region of high similarity.

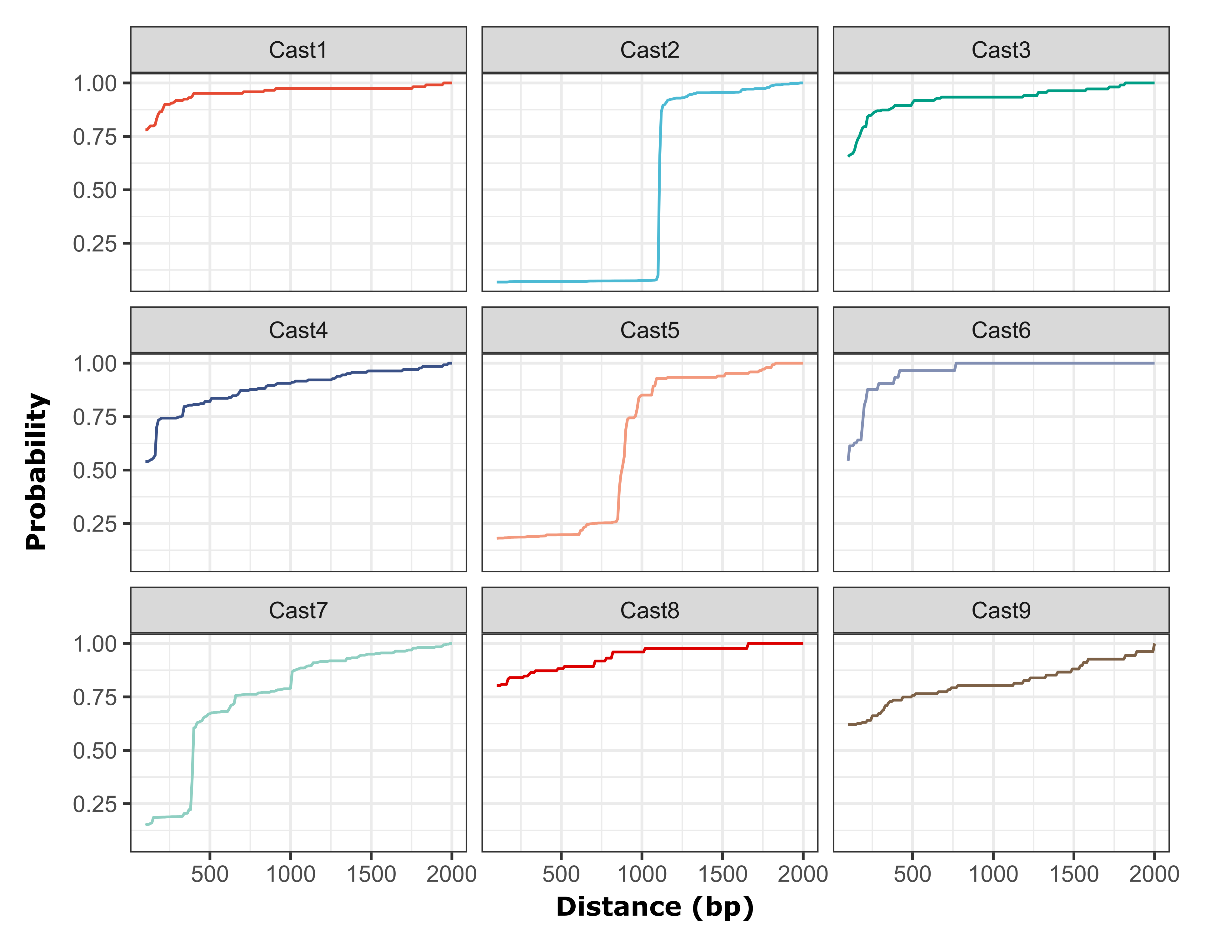

**Supplementary figure 4.** Analysis of the Cast1-Cast9 monomer organization. Probability represents the relative value of finding another Cast monomer at a set distance from another. Steep increases in Cast2, Cast5 and Cast7 indicate the presence of a particular pattern of satDNA organization that includes additional sequences. The rest of satDNAs do not have any sharp increases showing a tandem organization consisting only of satellite monomers with no additional sequences, evident from the relatively slow increases in the graph and a high starting point. As an example, Cast3 shows that 60% of all monomers in the genome are found immediately following another monomer. Cast2 has steep increase due to forming a new repeat unit Cast2’ (the increase is due to interCast2’ from Suppl figure 5.), Cast5 is often intercalated with R66-like elements and Cast7 comes in some instances with low similarity blast scores with TCAST but there not present in proper tandem organization.

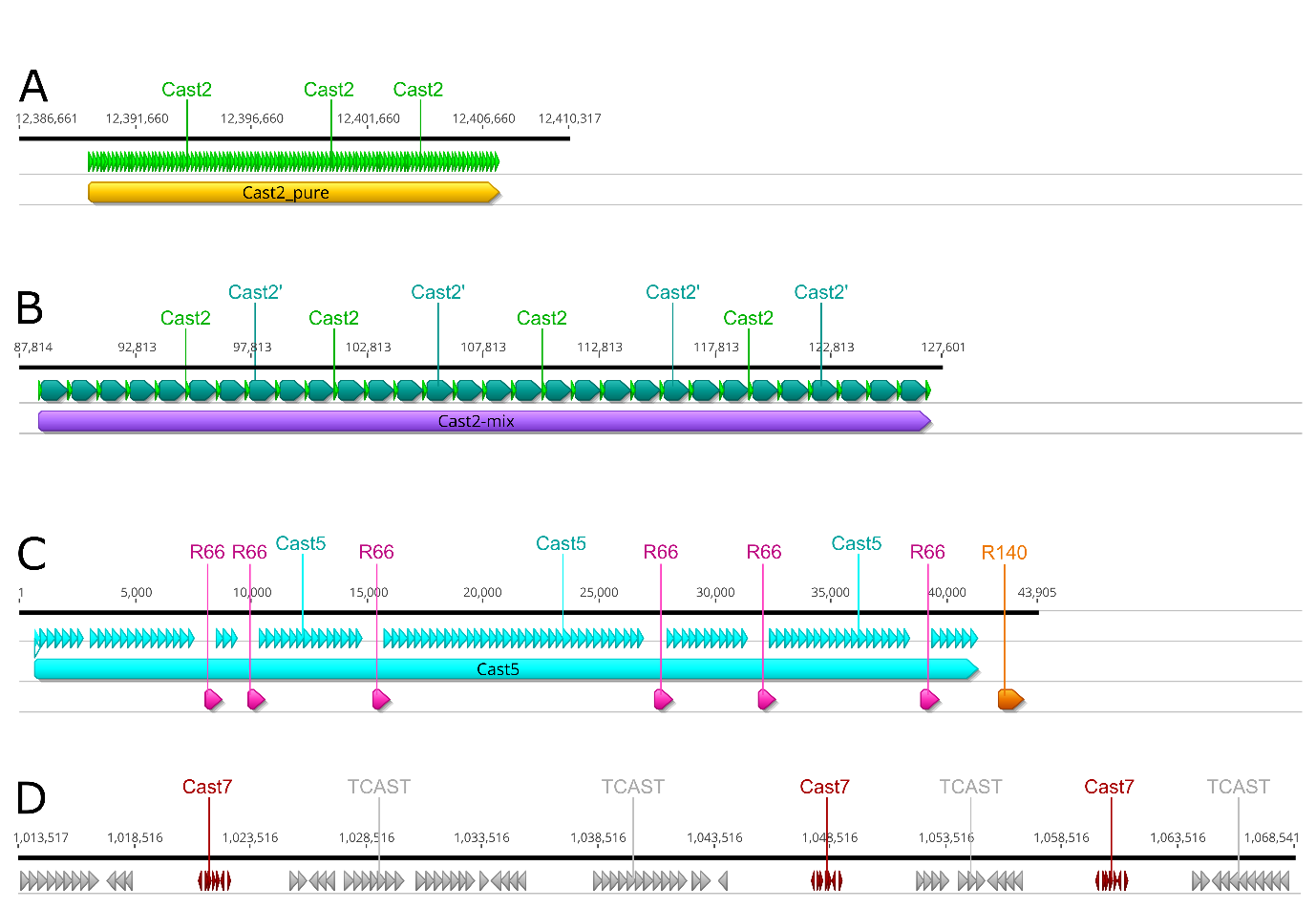

**Supplementary figure 5.** **A** Cast2 arrays in classical tandem head-tail organization. **B** Cast2 monomers (light green) with intermediate sequence (dark green) make up a new satDNA family Cast2’. **C** Cast5 arrays with adjacent regions (R66 and R140). **D** Cast7 arrays in association with (peri)centromeric TCAST satDNA.

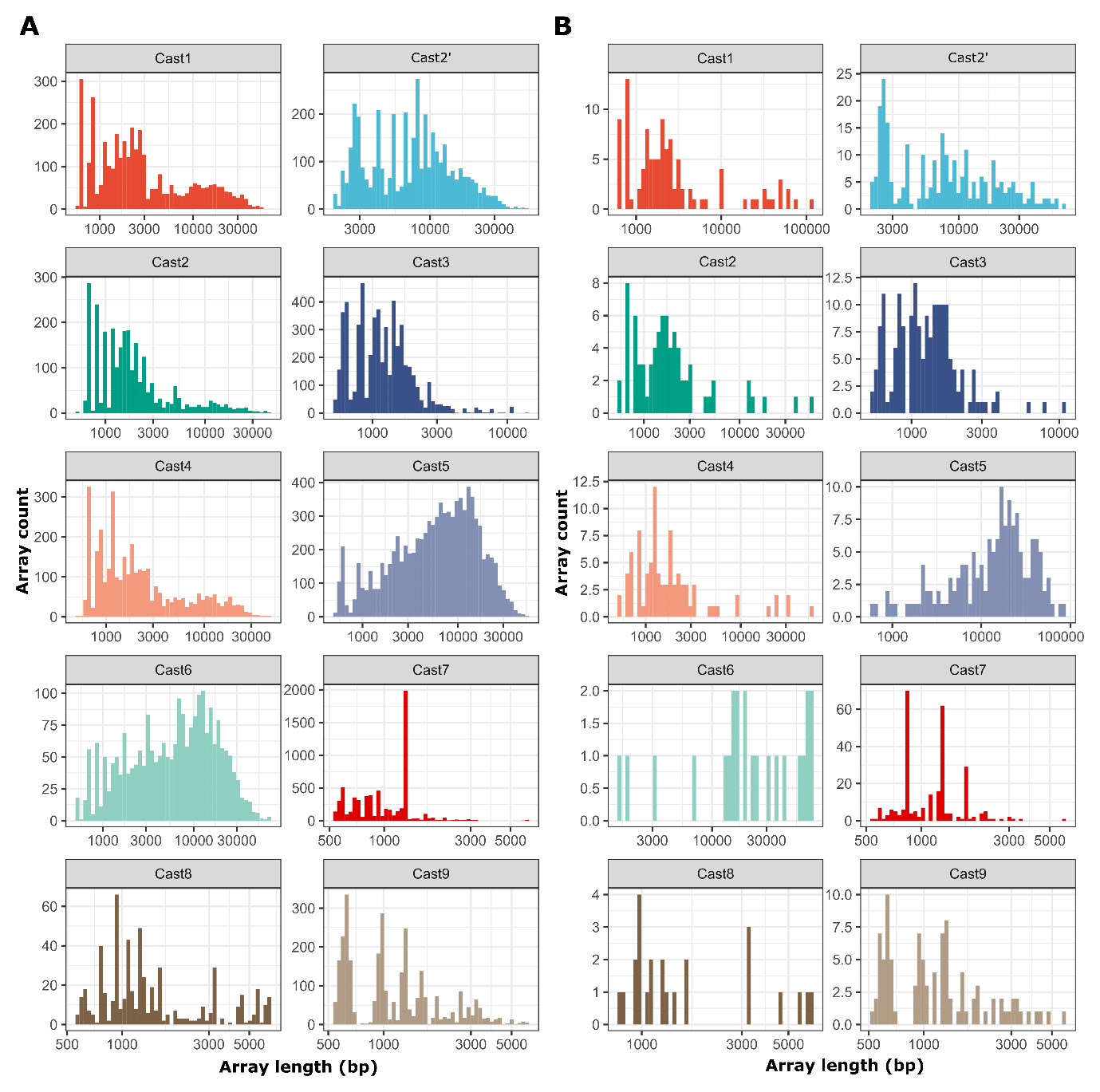

**Supplementary figure 6.** Graphical representation of individual Cast array lengths. **A** Cast arrays found in the longest raw Nanopore reads (50x coverage). **B** Cast arrays present in the TcasONT genome assembly. Array lengths shown on the x-axis are log scaled to better determine the spread of array lengths in both the genome and reads. The individual counts are separated based on different monomers and standardized based on total read coverage of the genome

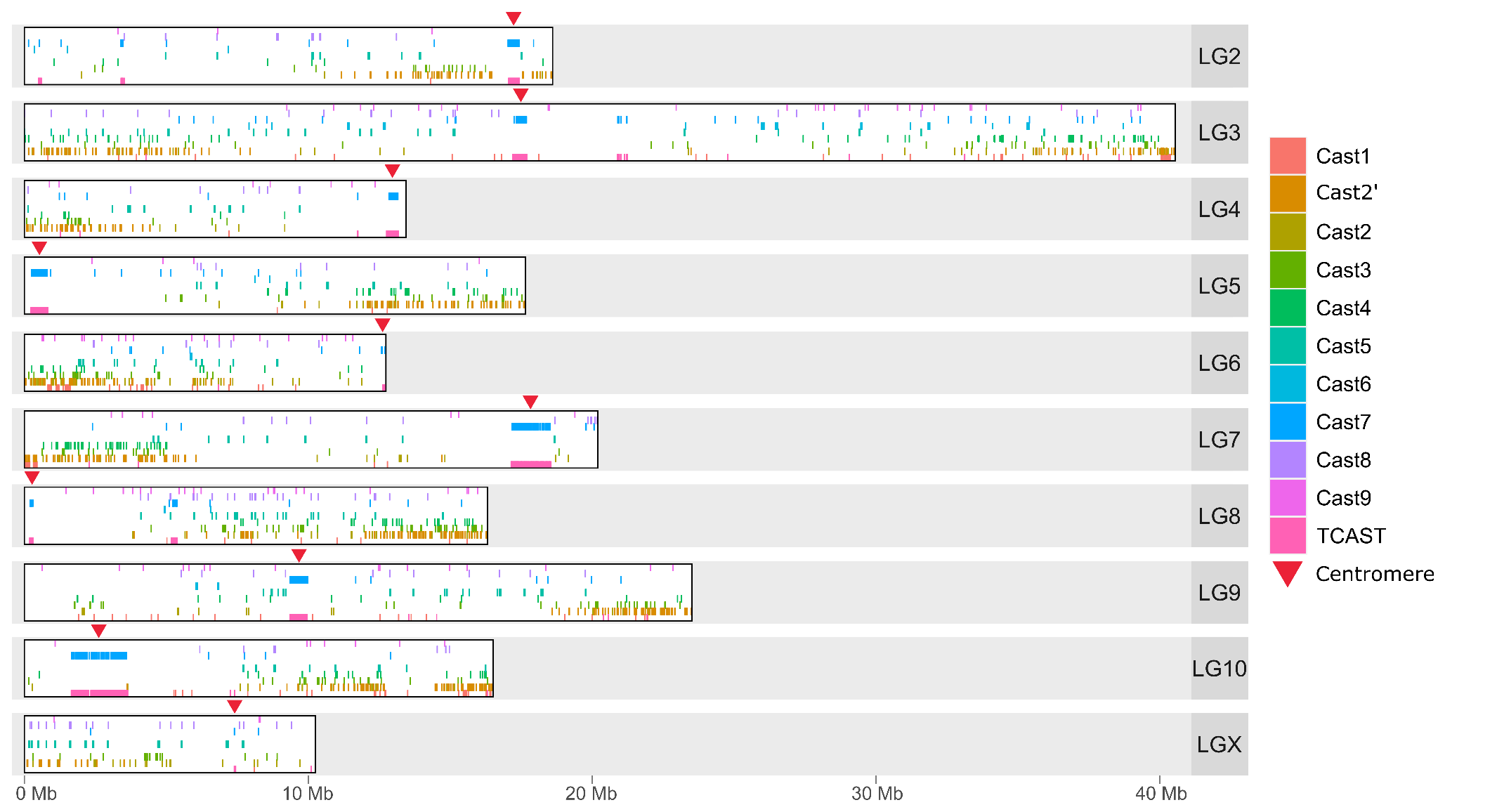

**Supplementary figure 7.** Localization of the Cast1-Cast9 satDNAs and TCAST on 10 linkage groups (chromosomes) of the TcasONT genome assembly. Location of centromere on each chromosome is indicated by a red arrow.

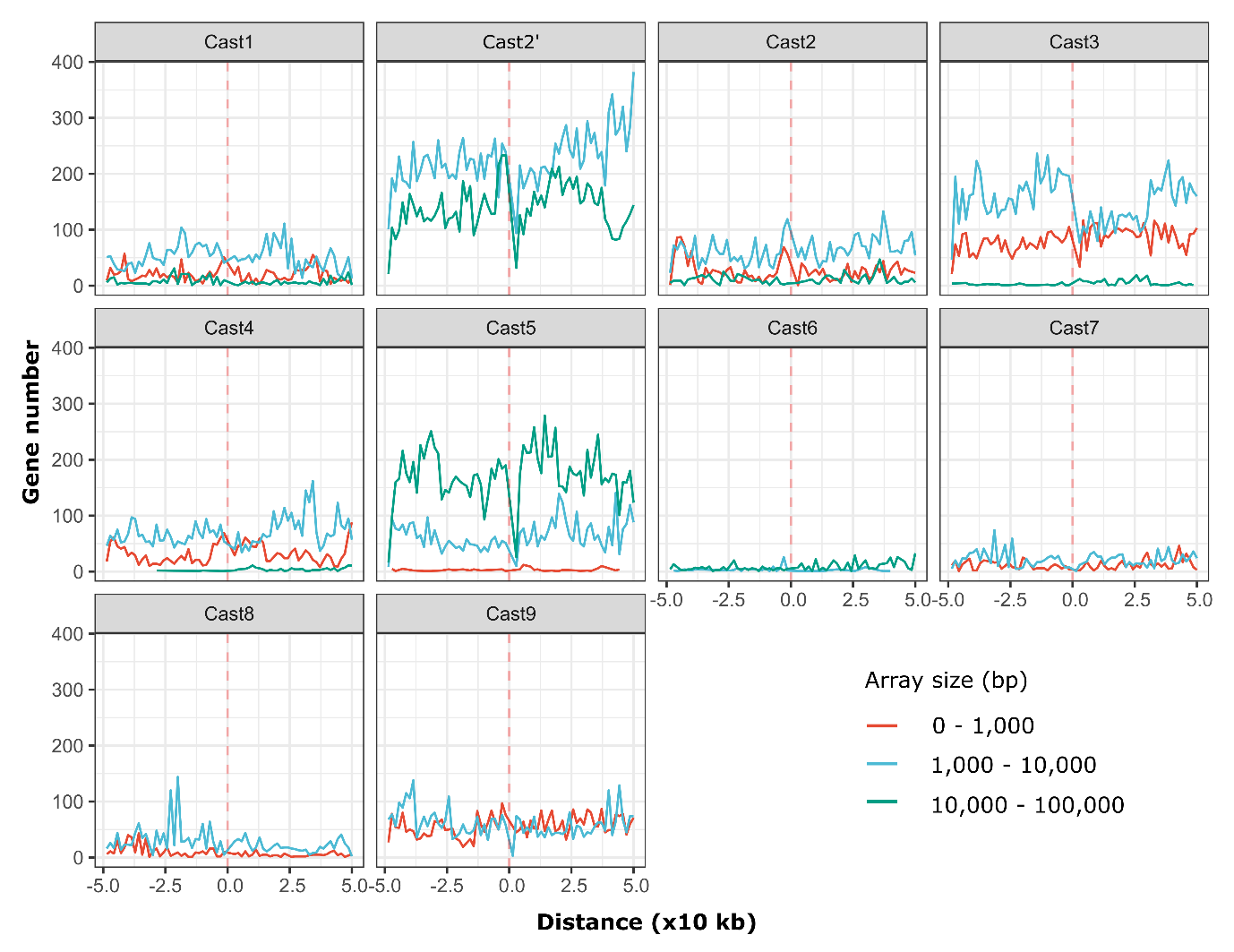

**Supplementary figure 8.** Relationship of Cast1-Cast9 satDNAs’ arrays and the number of genes (exons) in their vicinity. SatDNAs’ arrays are divided based on their length into three classes: short (<1000 bp), medium (1-10 kb) and long (10-100 kb). Counts represent the total sum of genes present in 1 kb bins 50 kb upstream and downstream of a specific Cast array, respectively.

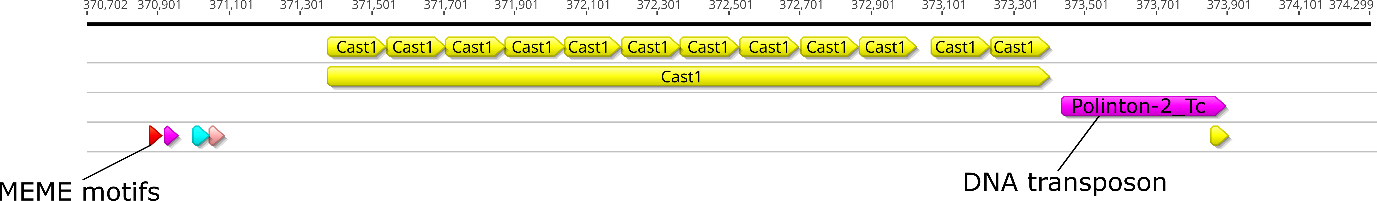

**Supplementary figure 9.** Pattern of Cast1 arrays with transposon-like sequence on one side of surrounding regions characteristic of all Cast1 arrays on chromosome 7 (LG7).

**A**

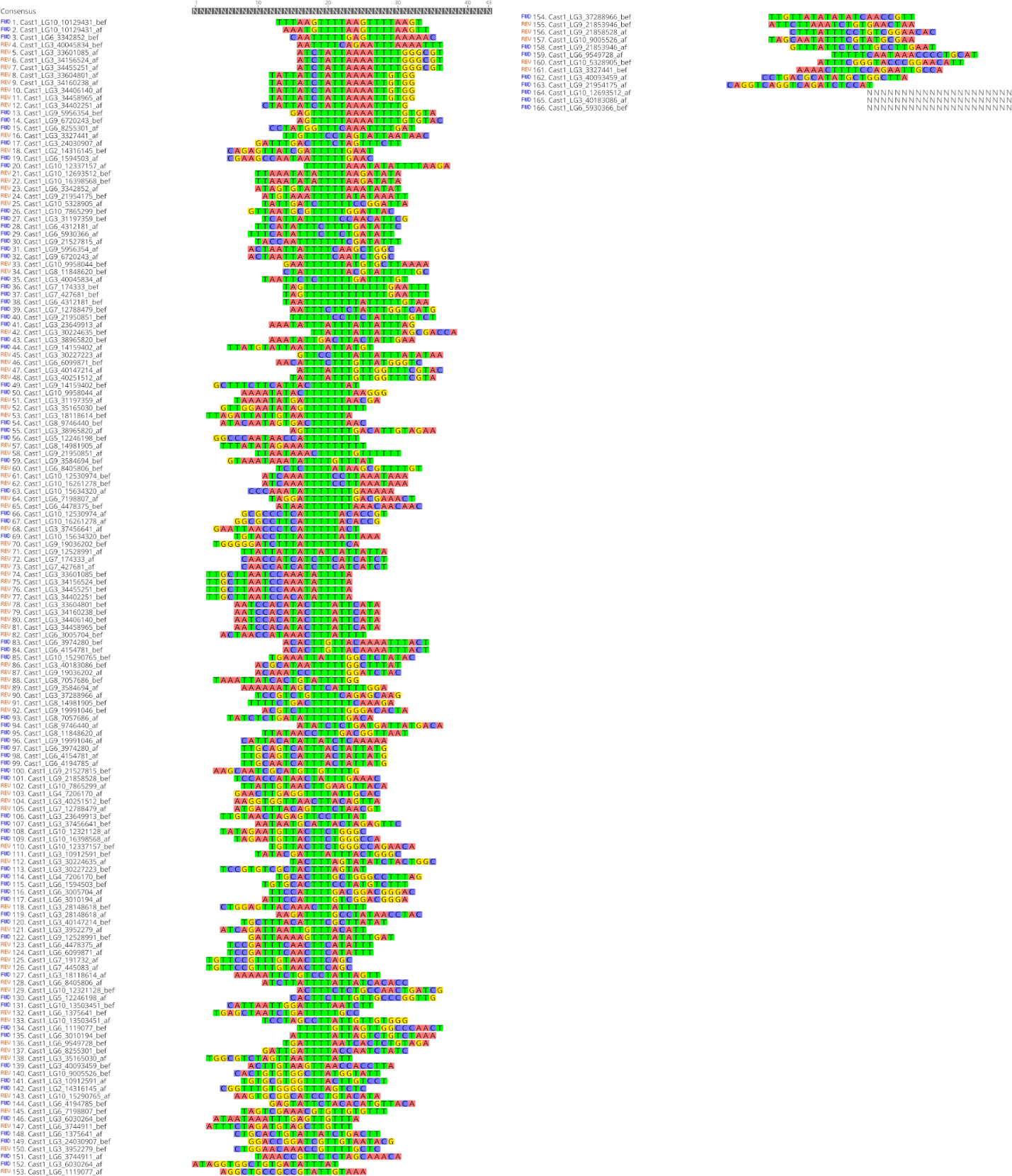

**B**

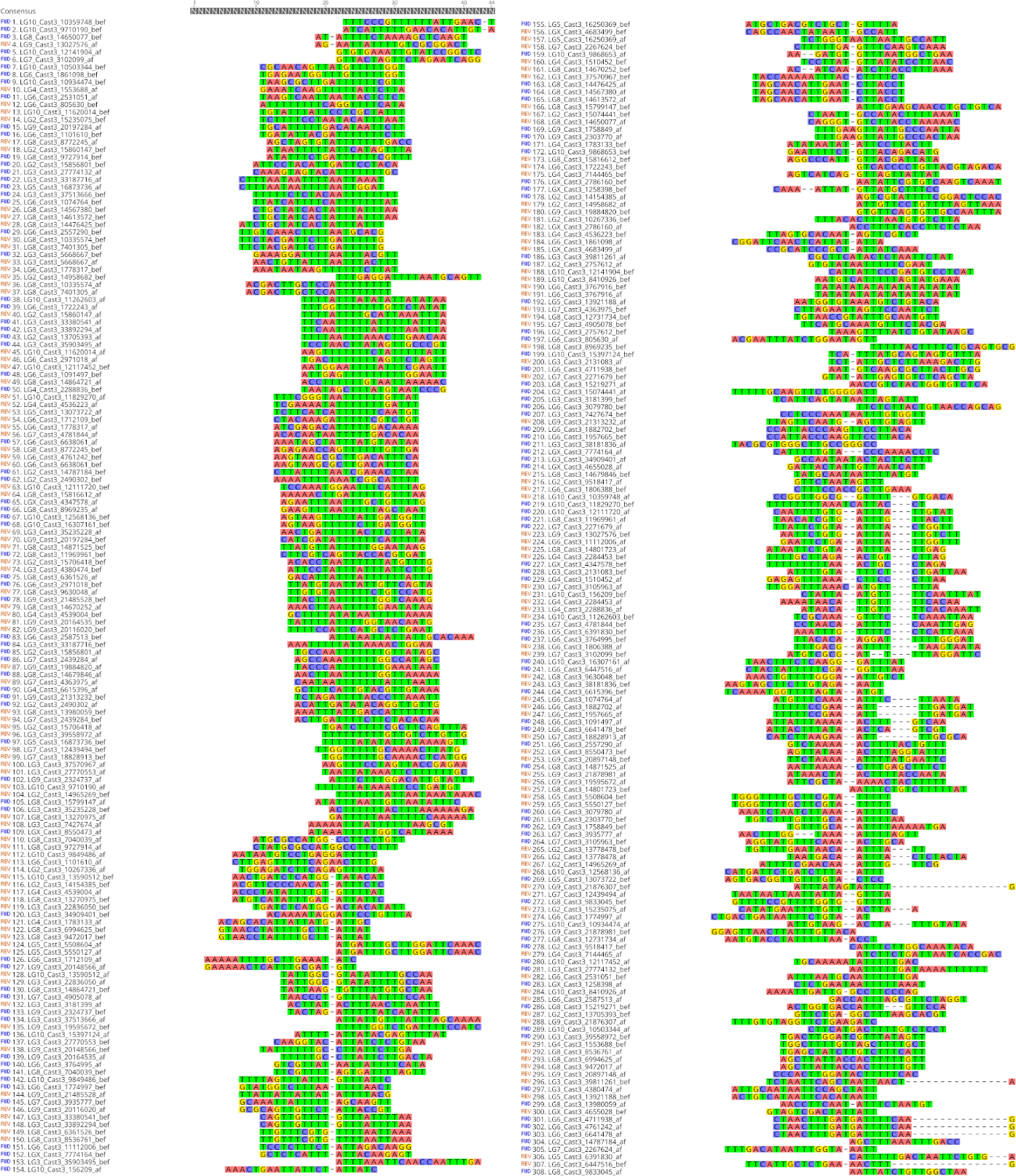

**C**

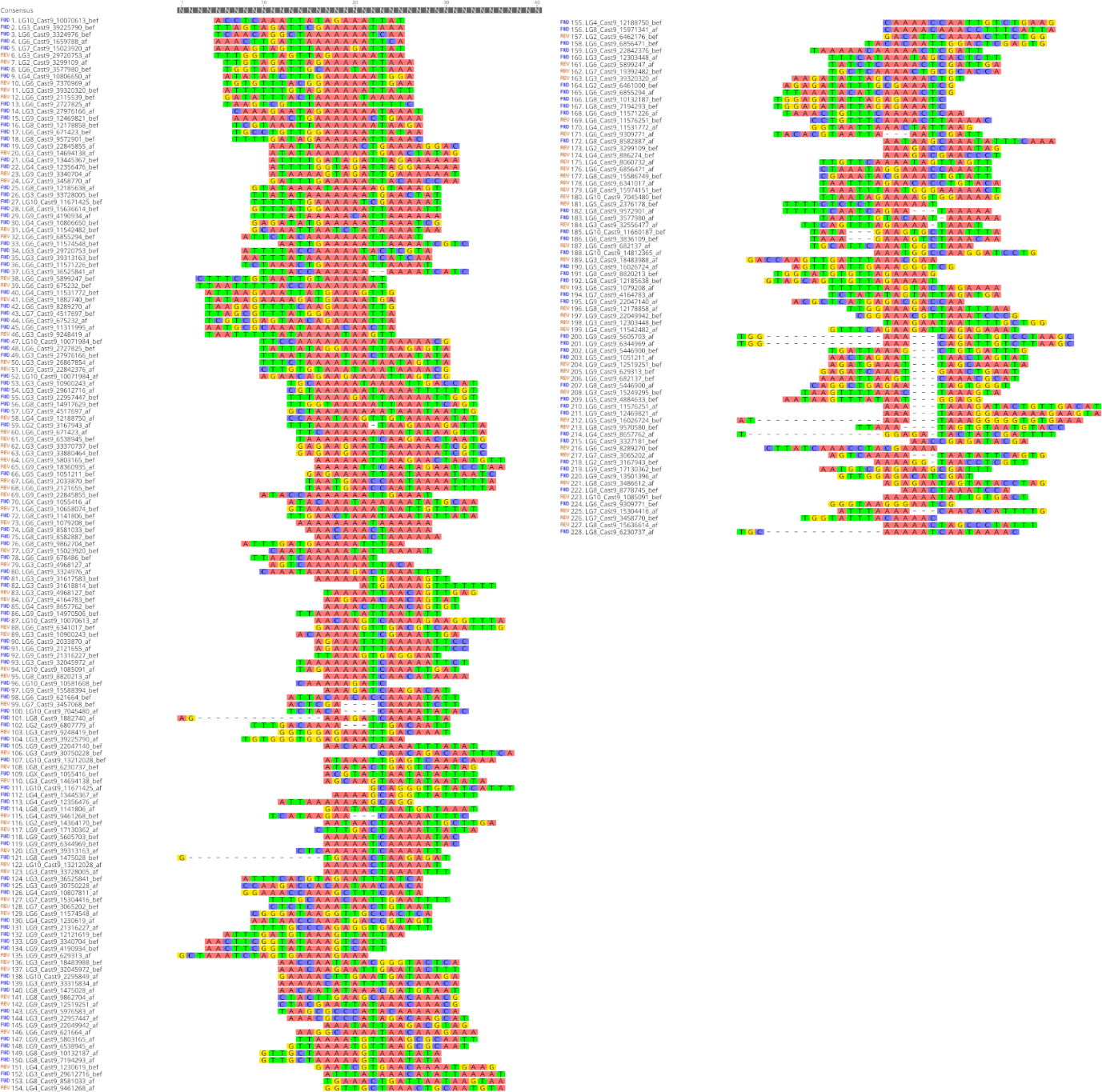

**Supplementary figure 10.** Junction region alignment for **A** Cast1 **B** Cast3 and **C** Cast9 where enrichment of polyT and polyA stretches is visible. Sequence motif are shown in Fig 5D. Precise edges were determined using the outlined k-mer counting algorithm and then sequences consisting of 20 bp before and after each precisely defined array edges were extracted and aligned using MAFFT algorithm.

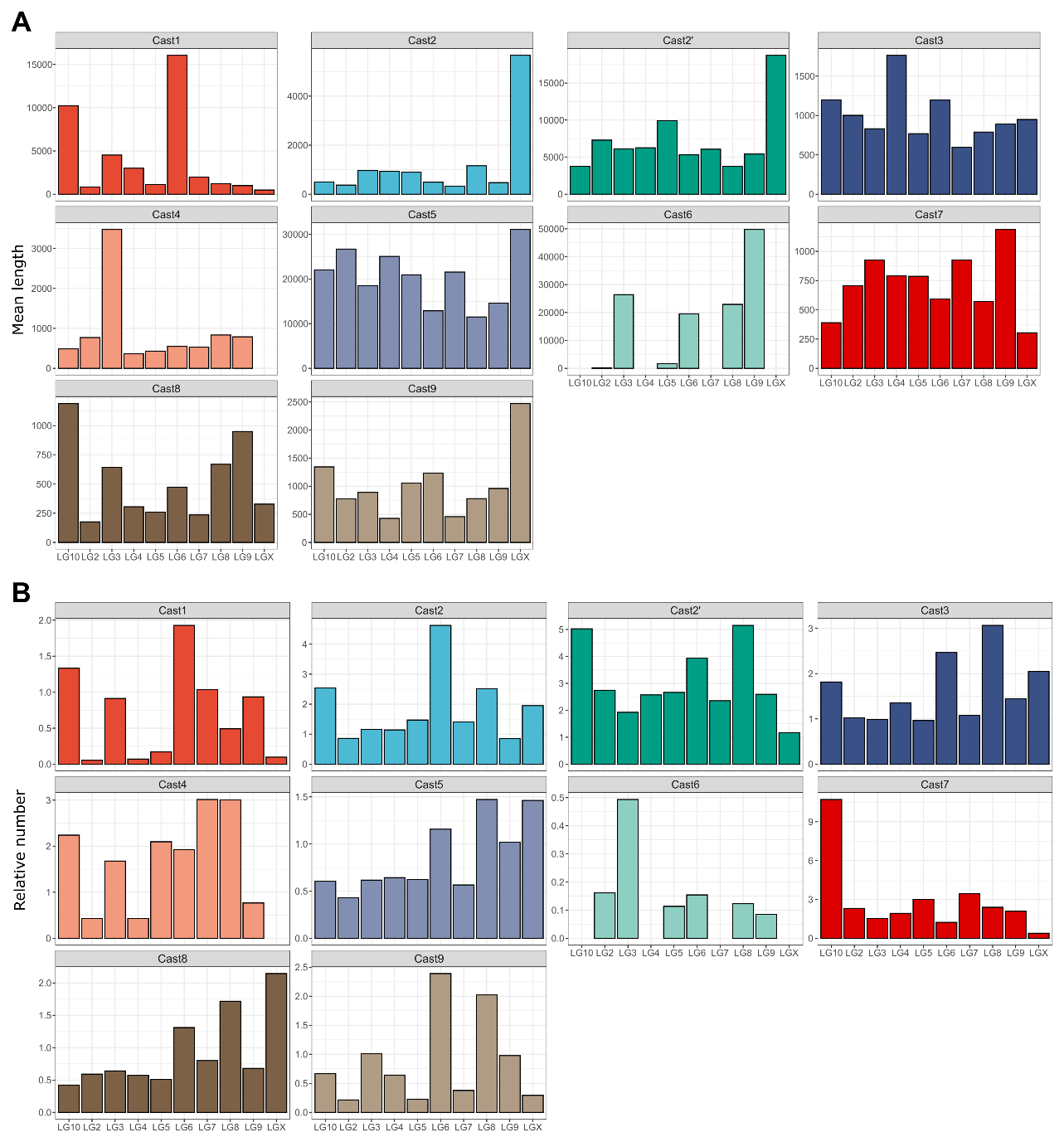

**Supplementary Figure 11. A** Mean length of arrays for each Cast on 10 different chromosomes (where Cast is present). **B** Standardized number of arrays per megabase for each of the chromosomes. The most prominent feature of elongation on X chromosome is visible for Cast2, Cast2’ Cast5 and Cast9, all exhibiting a large increase in their mean length and little divergence in their relative number per megabase.
